# Genome sequence and efficient CRISPR/Cas9 gene editing in the solanacea specialist pest *Tetranychus evansi*

**DOI:** 10.64898/2026.02.23.707505

**Authors:** Dries Amezian, Femke De Graeve, Rohith Mettumpurath Sasi, Lotte Vanhaecht, Antonio Mocchetti, Ernesto Villacis Perez, Merijn R. Kant, Sander De Rouck, Thomas Van Leeuwen

## Abstract

*Tetranychus evansi* is an invasive spider mite pest of solanaceous crops worldwide. Its global expansion and ability to rapidly develop acaricide resistance highlight the need for robust genomic resources and functional genetic tools to design custom control strategies. Here, we deliver a high-quality genome assembly and establish efficient CRISPR/Cas9 editing in *T. evansi* to enable mechanistic studies of host adaptation and pesticide resistance. Using an inbred line and Oxford Nanopore Technologies (ONT) long-read sequencing (∼36x), we assembled an 89 Mb genome into 13 contigs (N50 = 18.6 Mb) with high completeness and annotated 14,246 protein-coding genes. We manually curated the principal detoxification gene families (P450s, CCEs, GSTs, UGTs, ABC transporters and DOGs), revealing consistently smaller repertoires than in the highly polyphagous relative *Tetranychus urticae*.

To enable reverse genetics, we adapted SYNCAS (saponin + branched amphiphilic peptide capsules) for maternal delivery of CRISPR/Cas9 in *T. evansi*. Targeting the *phytoene desaturase* (*PD*) pigmentation marker produced reliable knockouts with visible albinism and mean editing efficiencies of ∼9.6-14.7%, allowing establishment of stable mutant lines. We further applied precision gene editing to knock-in (KI) the M918T and M918L substitutions into the voltage-gated sodium channel (VGSC). While M918L was lethal in *T. evansi*, we generated multiple homozygous lines for M918T (mean KI ∼4.4%). Bioassays demonstrated that while the mutation caused extremely high levels of bifenthrin resistance (RR>1000), this was less so for cyfluthrin (RR=164), revealing the independent and specific role of M918T in pyrethroid resistance.

Collectively, these resources establish *T. evansi* as a tractable system for reverse genetic analysis and provide a reference for future comparative and population genomics.

## 1. Introduction

The tomato red spider mite, *Tetranychus evansi* (Baker & Pritchard), is a major pest of solanaceous crops such as potato (*Solanum tuberosum*), tobacco (*Nicotiana tabacum*) and especially tomato (*Solanum lycopersicum*) (Navajas et al., 2013). Over the last three decades, *T. evansi*, a species endemic to South America (Silva, 1954), has expanded its geographical range and emerged as a serious invasive agricultural pest worldwide, causing significant damage (Azandeme-Hounmalon et al., 2022; Boubou et al., 2011; Migeon et al., 2009; Migeon and Dorkeld, 2026). Recent records (Azandeme-Hounmalon et al., 2022; Fan et al., 2021; Knihinicki et al., 2025; Mocchetti et al., 2025b; Monjarás-Barrera and Sanchez-Peña, 2024) suggest further expansion accelerated by ongoing climate change, thereby increasing threat to temperate regions (Ghazy et al., 2019; Yan et al., 2024). Yet, genomic resources and genetic tools are still lacking for this species, hampering the investigation of genetic variation associated with host plant use and pesticide resistance, limiting the efficiency of pest control strategies.

Although *T. evansi* is generally considered a specialist on Solanaceae, it has been reported on more than 140 plant species across 38 families (Migeon and Dorkeld, 2026), mostly as sporadic and circumstantial occurrences (Navajas et al., 2013). However, an expansion of host range has been documented in the Mediterranean basin and Africa, where *T. evansi* colonizes a broader diversity of plant species (Azandémè-Hounmalon et al., 2015; EPPO, 2026; Navajas et al., 2013). This expansion highlights the need for comprehensive genomic resources to better assess the genetic basis underpinning the adaptive potential of *T. evansi* and to anticipate shifts in host exploitation.

Control of *T. evansi* reliesheavily on chemical pesticides, such as organophosphates, pyrethroids and METI acaricides (Azandeme-Hounmalon et al., 2022; Carvalho et al., 2012; De Rouck et al., 2023; Nyoni et al., 2011), which results in high levels of resistance (De Rouck et al., 2023). However, relatively few studies have investigated the molecular basis of resistance, partly due to the lack of genomic resources. Resistance to organophosphates was linked to a mutation in one of the two genes coding for the acetylcholinesterase enzyme (AChE), *ace-1*, at position 331 (Carvalho et al., 2012). A recent screen of molecular resistance markers in Vietnamese populations identified for the first time in *T. evansi* the I321T substitution in the glutamate-gated chloride channel 3 (GluCl3), and a peculiar histidine-to-glutamine substitution at position 92 in the PSST subunit, a position previously linked to resistance to abamectin and METI-I acaricides respectively (Alavijeh et al., 2020; Bajda et al., 2017; Mocchetti et al., 2025b; Xue et al., 2022). Resistance to pyrethroid insecticides and acaricides is often associated with point mutations in the *voltage-gated sodium channel* (*VGSC*) gene that reduce insecticide sensitivity, the so-called knockdown resistance (“*kdr*”) (Feyereisen et al., 2015; Rinkevich et al., 2013). The most widespread mutation, L1014F, confers only moderate levels of resistance and has been reported in numerous arthropod pest species, although it is infrequent in mite species, where alternative mutations, such as F1538I/L, L1024V and L925V, are considered to be the main contributors to resistance and are also commonly referred to as *kdr* (Ding et al., 2015; González-Cabrera et al., 2013; Millán-Leiva et al., 2021; Rameshgar et al., 2019; Wu et al., 2019). Higher resistance is typically associated with the combination of *kdr* mutations with M918T, termed *super-kdr* for “super knockdown resistance” (Rinkevich et al., 2013). Notably, Nyoni et al. (2011) reported the presence of M918T in *T. evansi* in the absence of any other *kdr*, representing the first documented case of this mutation occurring alone in arthropods. Resistant populations from Malawi carrying M918T alone displayed high levels of resistance to pyrethroids, suggesting that this mutation may confer substantial resistance without the canonical *kdr* background. This finding challenged the traditional model of *super-kdr* evolution and highlights *T. evansi* as a valuable system for investigating this alternative form of target-site resistance.

The availability of a high-quality genome for *T. evansi* is instrumental for implementing genetic mapping studies capable of unravelling genetic determinants of host plant adaptation and pesticide resistance. It also enables high-resolution comparison of genome-wide gene expression patterns across populations. In addition, it facilitates the development and validation of reverse genetic tools, such as CRISPR/Cas9 and RNA interference (RNAi), by providing the precision needed to identify gene function and verify potential off-target effects. Understanding the molecular and genetic mechanisms underlying the development of resistance is key for developing adequate pest control strategies. When specific genes or mutations are associated with resistance, CRISPR/Cas9-based genome editing can be a powerful tool to validate the actual contribution of these genetic factors to the observed resistance (Douris et al., 2020). Recently, the development of maternal delivery of CRISPR/Cas9 components has emerged as an effective alternative to the traditional embryo injection (De Rouck et al., 2024; Li et al., 2026, 2021; Mocchetti et al., 2025a; Shirai et al., 2022). The SYNCAS approach, initially developed for *Tetranychus urticae*, exploits the synergistic effect of saponins and branched amphiphilic peptide capsules (BAPC) in a formulation that enhances delivery efficiency (De Rouck et al., 2024). Since its development, SYNCAS has been successfully applied in various insect and mite species (Guerra et al., 2026; Matsuda et al., 2025; Mocchetti et al., 2025a). It was used to validate resistance mechanisms toward METI-II acaricides in spider mites (İnak et al., 2024a), spinosad resistance in thrips (Mocchetti et al., 2025c) and amitraz resistance in *Varroa* mites (İnak et al., 2024b). It also demonstrated that loss-of-function in the nAChR α6 subunit confers resistance to spinosyns in spider mites and predatory mites, confirming mode of action in chelicerates (Mocchetti et al., 2026).

Here, using Oxford Nanopore Technologies (ONT) long-read sequencing, we provide a fully annotated genome with extensive manual curation of the main detoxification gene families. We also develop a gene-editing approach by targeting a visible marker of pigmentation, a *phytoene desaturase* (*PD*) gene. We then further use SYNCAS-based precision editing to functionally validate the specific and independent role of M918T in the *VGSC* associated with pyrethroid resistance in *T. evansi*.

## 2. Materials and Methods

### 2.1. Mite rearing and inbreeding

The genome of *T. evansi* was assembled from sequencing an inbred line derived from the Viçosa-1 strain, collected in Brazil in 2002 (Sarmento et al., 2011) and maintained in the lab on detached leaves from tomato plants (*Solanum lycopersicum* var. “Money Maker”), without acaricide treatment. Viçosa-1 was inbred via mother-son mating performed on detached tomato leaves as described by Bryon et al. (2017a) for 7 generations to generate the inbred strain (hereafter suffixed with the “i” for inbred designation: Viçosa-1i). After the final round of inbreeding, Viçosa-1i (VICi) was expanded and maintained on potted tomato plants to produce sufficient mites for downstream analyses. A second *T. evansi* strain, bean-adapted (TEBA), generated at the University of Amsterdam and used for gene-editing experiments for practical reasons. To confirm species identity, *COI* markers from TEBA and VICi were amplified with primers and conditions described in Mocchetti et al. (2025b), and compared to sequences from various spider mite populations (Supplementary Figure 1, Mocchetti et al., 2025b). The TEBA strain was maintained on potted bean plants (*Phaseolus vulgaris* cv. “Prelude”). All strains were maintained at 25 °C (± 0.5°C), 60% relative humidity, and a 16:8 h light:dark photoperiod.

### 2.2. Genome assembly

#### 2.2.1. DNA and RNA extraction, library prep and sequencing

High-molecular-weight genomic DNA was extracted from more than 10,000 adult mites. Homogenization was performed in 950 µL of SDS buffer (200 mM Tris-HCl, 400 mM NaCl, 10 mM EDTA, 2% SDS, pH 8.2), 18 µL of Proteinase K (10 mg/mL) and 3.6 µL of RNase A (40mg/mL). Samples were incubated at 60 °C with constant agitation for 2 h, followed by an additional incubation of 1 h 30 min at 37 °C after the addition of 3.6 µL RNase A. Subsequently, a standard phenol-chloroform extraction was performed (Van Leeuwen et al., 2008). DNA concentration and quality were assessed using a Qubit Fluorometer with the Qubit dsDNA HS Assay kit (Thermo Fisher Scientific, USA) and the 4200 Tapestation System with the Genomic DNA ScreenTape assay (Agilent), respectively. Sequencing libraries were prepared following the manufacturer’s recommendations for ONT sequencing and sequenced on an R10.4 flow cell by Ohmx.bio (Gent, Belgium).

To support genome annotation, total RNA was extracted from a pool of 400 individuals across different developmental stages using the RNeasy Plus Mini Kit (Qiagen, Germany) following the manufacturer’s protocol. RNA concentration and integrity were assessed using a DeNovix DS-11 spectrophotometer (DeNovix, USA) and agarose gel electrophoresis (1% agarose, 30 min, 100 V). Directional libraries were prepared by Novogene (UK) and sequenced on an Illumina NovaSeq X platform to generate 150-bp paired-end reads.

#### 2.2.2. Assembly

Raw reads were first trimmed with Seqtk (v1.3-r106) to remove 50 bp from both read ends and reads containing adapter sequences were filtered out using Seqkit (v2.0.0). To improve assembly contiguity, only reads with a minimum length of 10 kb were retained for downstream processing. The filtered long reads were assembled with Flye (v2.9.3) using an expected genome size of 90 Mb, with default parameters unless otherwise specified. The draft assembly was subsequently polished with Medaka (v2.0.1), which corrects systematic base-calling errors using the original ONT long-read data. Putative mitochondrial contigs were identified based on sequence similarity to known arthropod mitochondrial genomes and removed from the assembly to avoid redundancy with the nuclear genome assembly. Assembly completeness was evaluated with BUSCO (v5.4.6) in genome mode using the *arachnida_odb10* database. Repetitive elements were identified *de novo* with RepeatModeler (v2.0.6) and masked with RepeatMasker (v4.1.9) in soft-masking mode to prepare the assembly for annotation.

#### 2.2.3. Genome annotation

Structural and functional annotation of the *T. evansi* genome was carried out with Funannotate (v1.8.17). The annotation pipeline integrated multiple sources of evidence including RNA-seq data, transcript assemblies, and protein databases. Protein sequences from *T. urticae* (Ji et al., 2023; Wybouw et al., 2019a) were aligned to the repeat-masked *T. evansi* genome assembly with Miniprot (v0.12-r237), and the resulting GFF3 file was incorporated as additional protein evidence into Funannotate with a weight of 10. The functional annotation of predicted genes was carried out using integrated databases and tools within Funannotate, including InterProScan, Pfam, EggNOG, and SignalP, to assign putative functions to gene models. In addition, reciprocal BLASTp searches (E-value threshold: 1 x 10^-5^) were performed against the *T. urticae* protein sequences. For each *T. evansi* gene with a reciprocal best BLAST hit to *T. urticae* (*i.e.* the top hit in both directions), the corresponding *T. urticae* gene description was assigned to the *T. evansi* gene, otherwise it was annotated as a hypothetical protein. The annotation was further processed to retain only the longest isoform per gene, and final NCBI-compatible submission files were generated with table2asn (v1.29.324).

Detoxification gene families were manually curated to improve annotation accuracy. Cytochrome P450 genes (P450s) were identified based on the presence of the InterPro domain IPR001128, and each predicted sequence was queried using tBLASTn against the genome and BLASTp against a curated P450 database (Arthropod P450 enchiridion; arthropodp450.eu (Charamis et al., 2025)) to confirm gene structure. Gene length discrepancies or exon structure issues were corrected when supported by the sequence evidence, and adjacent fragments were merged when they clearly represented a single gene. To detect exons or gene copies that might have been missed by the annotation, extended genomic regions around problematic loci were searched using BLASTx against the P450 database. Fragments that could not be reliably reconstructed (e.g. isolated exons) were excluded.

For the remaining detoxification gene families, carboxyl/choline esterases (CCEs), glutathione S-transferases (GSTs), UDP-glycosyltransferases (UGTs), intradiol ring-cleavage dioxygenases (DOGs), and ATP-binding cassette (ABC) transporters, candidate genes were initially identified based on the presence of family-specific InterPro domains (CCEs: IPR002018; GSTs: IPR004045; UGTs: IPR002213, IPR050271; DOGs: IPR015889; ABCs: IPR003439). Predicted gene models were subsequently manually inspected and adjusted as needed, including correction of exon boundaries, merging of split models when supported by sequence and phylogenetic evidence, and exclusion of incomplete and misannotated predictions based on sequence evidence.

Phylogenetic analyses were conducted to validate gene family assignments, identify potentially misannotated or chimeric gene models, and determine the clade-level placement of detoxification genes. Protein sequences were aligned using MAFFT (v7.526) under default settings, and maximum-likelihood trees were inferred using IQ-TREE (v3) with automatic substitution-model selection and 1,000 ultrafast bootstrap replicates (-m MFP -bb 1,000)(Wong et al., 2025). Phylogenetic trees were visualized and inspected using iTOL (Interactive Tree Of Life, Letunic and Bork, 2024).

#### 2.2.4. Inbred line heterozygosity

To compare heterozygosity profiles between the outbred and inbred Viçosa-1 strains, we performed SNP calling on whole genome sequencing (WGS) data following GATK’s (v4.2.0.0) (McKenna et al., 2010) best practices. Total DNA was extracted from 800 adult females per strain using the procedure described in Section 2.2.1. Reads were aligned to the *de novo* genome assembly using BWA-MEM (v0.7.17-r1188) with default settings, and duplicate reads were marked with Picard *MarkDuplicates*. Variants were called using GATK *HaplotypeCaller* and jointly genotyped across both strains, yielding a single VCF file for downstream analyses. Heterozygosity was quantified by counting heterozygous SNPs, defined as SNPs with an alternate allele frequency between 0.05 and 0.95 (values outside this range were considered fixed for either the reference or alternate allele), in sliding windows of 100 kb with a step size of 5 kb along the genome. Analyses were restricted to contigs ≥ 500 kb.

### 2.3. Development of CRISPR/Cas9 gene editing in *T. evansi*

#### 2.3.1. Target-genes and phylogenetic analysis

To test whether the CRISPR/Cas9-based SYNCAS formulation (De Rouck et al., 2024) can also be developed in to a gene editing tool for *T. evansi*, knockout (KO) experiments were performed targeting the *phytoene desaturase* (*PD*) gene, a genetic marker with a clear visible phenotype in spider mites (Bryon et al., 2017a; De Rouck et al., 2024; Dermauw et al., 2020a). Orthologues of *T. evansi* carotenoid synthase/cyclase (*CSC*) and desaturase genes were identified using a local tBLASTn (BLAST+, v2.16.0+) (Camacho et al., 2009) search against the newly assembled *T. evansi* genome, with *T. urticae* protein sequences as queries (*CSC* genes: *tetur01g11260*, *tetur11g04840; PD* genes: *tetur01g11270, tetur11g04810, tetur11g04820, tetur11g04830*; sequences available with the last version of the reference genome (Ji et al., 2023; Wybouw et al., 2019b)). Briefly, tBLASTn searches were run with an E-value threshold of 1 x 10^-5^. The candidate proteins were subjected to reciprocal BLASTp searches against the *T. urticae* proteome using identical parameters. The relative homology of *PD* and *CSC* genes with *T. urticae* sequences and representative sequences from arthropods, fungi, cyanobacteria, algae, and plants was further illustrated in a maximum-likelihood phylogenetic tree (Supplementary Table 2). Protein sequences were aligned with MAFFT (v7.526) under default setting and the resulting alignment was used to build a tree with IQ-TREE (v3.0.1, -m MFP -bb 1000)(Wong et al., 2025). The layout and decoration of the consensus tree were edited using ITOL (Letunic and Bork, 2024). Based on this analysis, the *PD* gene *TETEV_003852* was selected as target for CRISPR/Cas9 knockout.

A sgRNA targeting the *voltage-gated sodium channel* (*VGSC*; *TETEV_013537*) was also designed for validation of the M918T and M918L mutations, potentially responsible for pyrethroid resistance in *T. evansi* (Nyoni et al., 2011) and other arthropods (Rinkevich et al., 2013), using homology-directed repair (HDR)-mediated gene editing. The sgRNA target sequences were designed using the online tool CRISPOR (Concordet and Haeussler, 2018), where the guide with highest predicted efficiency (sgRNA *PD*) or closest proximity to the M918 mutation site (sgRNA *VGSC*) was selected. Repair templates used to knock in M918T and M918L were manually designed to introduce the corresponding codon substitution, along with additional silent mutations to further disrupt the target-site of the sgRNA, thereby preventing further Cas9 cleavage after HDR. Homology arms of 80bp in length were added on both sides flanking codon 918. Sequence information for sgRNAs and ssODN templates is provided in Supplementary Table 3.

#### 2.3.2. CRISPR-Cas9 injection mix

The SYNCAS formulation (De Rouck et al., 2024), with a few modifications, was used for the microinjections. Recombinant Streptococcus pyogenes Cas9 protein (Alt-R^Ⓡ^ S.p. Cas9 Nuclease V3, glycerol free, #10007808) was concentrated to 50 µg/µl using an Amicon® Ultra Centrifugal Filter (50 kDa MWCO, Merck Millipore), sgRNAs (Alt-R™ CRISPR-Cas9 sgRNA) and ssODN repair templates (Alt-R™ HDR Donor Oligo) were purchased from Integrated DNA Technologies. Lyophilized sgRNAs were dissolved at a concentration of 20 μg/μL in TE buffer. Repair templates were dissolved in nuclease-free water at a concentration of 1,000 μM. Branched Amphiphilic Peptide Capsules (BAPC) were purchased from Phoreus Biotech whereas saponin was obtained from Sigma Aldrich (cat#558255) and were dissolved in nuclease-free water at concentrations of 50 μg/μL and 1.3 μg/μL, respectively. The *PD* KO CRISPR/Cas9 injection mix was prepared by combining 1.5 μL Cas9 (50 μg/μL), 2 μL sgRNA *PD* (20 μg/μL), 0.5 μL BAPC (50 μg/μL), and 0.2 μL saponin (1.3 μg/μL), while the *VGSC* CRISPR/Cas9 mix was prepared with 1.8 μL Cas9 (50 μg/μL), 1.8 μL sgRNA *VGSC* (20 μg/μL), 0.5 μL ssODN (1,000 μM), 0.5 μL BAPC (50 μg/μL), and 0.2 μL saponin (1.3 μg/μL).

#### 2.3.3. T. evansi microinjections

CRISPR/Cas9 microinjections were performed as described previously (De Rouck et al., 2024). Since *T. evansi* is an arrhenotokous species (unfertilized eggs develop in haploid males), recessive mutations, such as a *PD* knockout, are visible in the phenotype of male offspring. For each replicate, about 400 females, 3-4 days in adult stage, were aligned on 2% agarose gel and injected ventrally in the body cavity between the third pair of legs with 3 nL of the CRISPR mix using a Nanoject III (Drummond Scientific). After injection, females were placed on detached bean leaves (on wet cotton) and allowed to lay eggs, with transfer to a fresh bean leaf every 24 h for 48 h, after which the injected mites were discarded. For *PD* KO experiments, three independent injections were performed, while a single injection set was conducted for KI of the M918L and M918T mutations.

#### 2.3.4. Screening and analysis of PD mutant mites

Males were crushed individually in 9 μL STE buffer (100 mM NaCl, 10 mM Tris-HCl, 1 mM EDTA, pH 8.0), and 1 μL Proteinase K (10 mg/mL). Females were crushed individually in 18 μL STE buffer and 2 μL Proteinase K. This crude extract was incubated at 37 °C for 30 min followed by Proteinase K deactivation at 98 °C for 10 min. A PCR was performed on these raw DNA extracts, amplifying a 516-bp region containing the sgRNA target site using GoTaq^Ⓡ^ G2 DNA Polymerase and the PD_F and PD_R primer pair (Supplementary Table 3). The temperature profile consisted of a 2-min denaturation step at 92 °C, followed by 40 cycles of 30 s at 92 °C, 30 s at 54 °C, and 1 min at 72 °C. After a final elongation of 72 °C for 5 min, a quality check was performed by running 5 μL of PCR product on a 2% agarose gel for 30 min at 100 V. The remaining PCR product was purified using an EZNA^Ⓡ^ Cycle Pure Kit (Omega Bio-Tek) before samples were sent to Eurofins Genomics (Germany) for Sanger sequencing.

#### 2.3.5. Establishment of VGSC M918L and M918T homozygous lines

Offspring of injected females were designated as the G0 generation. The haplodiploid sex determination of *T. evansi* facilitated directed crosses and the establishment of lines homozygous for the target mutations. Approximately 60-70 single G0 pairs were isolated on individual leaf discs and allowed to mate for two days (G1 crosses). Subsequently, both parents were processed for DNA sequencing (Section 2.3.3); males were crushed immediately after mating (*ca.* two days), while females were allowed to oviposit for up to six days before sequencing.

As no G1 pairs were recovered in which both individuals carried the desired M918T (or M918L, see Section 3.5) mutation, either in homozygous or hemizygous state, a second round of crosses was performed (G2 crosses). Specifically, 10-15 pairs were established using female progeny of a G1 mutant male, and male offspring of a G1 mutant female. In this scenario, all females used in G2 crosses were heterozygous for the M918 mutation, while 50% of the haploid males were expected to be mutants. After a two-day mating period, all G2 males were sequenced. In cases where the male was a confirmed mutant, the resulting female offspring were isolated individually on single leaf discs, kept unfertilized, and allowed to oviposit for up to three days. Finally, each female was backcrossed to one of her sons, allowed to lay eggs for up to six days, before being sequenced. Through this crossing scheme, four homozygous lines carrying the M918T mutation were established.

### 2.4. Chemicals and bioassays

Commercial formulations of all pyrethroid insecticides – type-I pyrethroid bifenthrin (Talstar 8 SC), and type-II pyrethroid beta-cyfluthrin (Bulldock 25 EC) – were used for toxicity assays. Dose-response assays were conducted with adult female mites as previously described for *T. urticae* (Khajehali et al., 2011). Briefly, 20-25 adult females were placed on the upper side of square kidney bean leaf discs placed on wet cotton. A custom-built spray tower was used to spray a total of 1 mL of acaricide solution at 1 bar pressure for each replicate (1.95 ± 0.05 mg acaricide deposited per cm^−2^). Mortality was assessed after 24 h, and all assays were performed with at least five acaricide doses and three replicates. Sprayed leaf discs were placed in a climate room with 26 ± 1 °C, 60 ± 2% RH, and a 16 L:8D photoperiod without direct light exposure. Mites that could not move when prodded with a fine brush under a stereomicroscope were considered dead. Control discs were only sprayed with deionized water, and mortality rates of control discs were always <10%. LC_50_ values and their 95% confidence intervals were calculated using POLOPlus-PC software (Robertson et al., 2017). LC_50_ values of WT and *VGSC^M918T^* TEBA mites for a given acaricide were considered significantly different if their 95% confidential intervals did not overlap (Robertson et al., 2017). Resistance Ratios (RR) were calculated by dividing LC_50_ values.

## 3. Results

### 3.1. Genome assembly and annotation

Starting with a single virgin female from the Viçosa-1 strain, we performed inbreeding with mother-son crosses for seven generations. The subsequent strain, thereafter denoted Viçosa-1i, was expanded on intact tomato plants and used for high molecular weight DNA extraction. Based on Oxford Nanopore Technologies (ONT) long-read sequencing (∼36x), the *T. evansi* genome was assembled into 13 contigs spanning 89 Mb, with a contig N50 of 18.6 Mb and a maximum contig length of 31.5 Mb. The assembly had a GC content of 31.9%. Two contigs likely corresponding to the mitochondrial genome (identified by BLAST) were excluded from the assembly. BUSCO analysis indicated 94.8% completeness against the *arachnida_odb10* dataset, including 89.5% single-copy and 5.3% duplicated genes. Repetitive elements accounted for 16.4% of the genome.

Variant calling for the Viçosa-1 outbred and related inbred strains identified 79,386 variants, of which 64,366 were SNPs. This relatively low number of variants is consistent with the use of the derived Viçosa-1 inbred line as the assembly source for the reference genome. In addition, the data shows that 7 generations of inbreeding were sufficient to achieve homozygosity at more than 99% of previously polymorphic sites, yielding a highly inbred Viçosa-1i strain (Figure 1).

**Figure 1:**
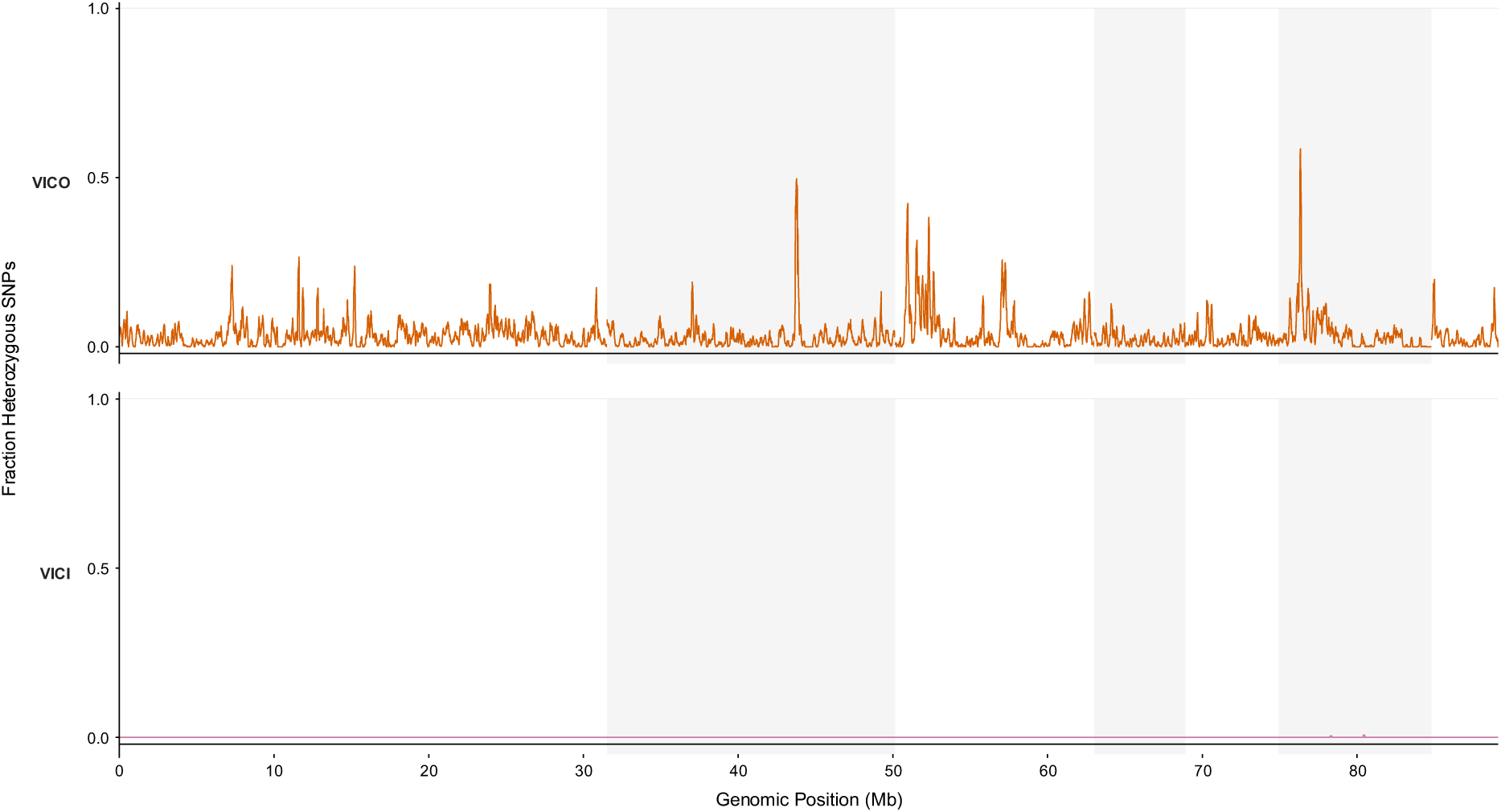
Genome-wide fraction of heterozygous SNPs in Viçosa-1 and Viçosa-1i. Genome-wide heterozygosity quantified by the fraction of significant heterozygous SNPs (FDR < 0.01) as assessed with sliding window analysis (100 kb window, 5 kb step-size). Contigs ( ≥ 500 kb) are arranged in descending order and indicated by alternating shading. Upper panel: outbred, Viçosa-1 (VICO); lower panel: inbred, Viçosa-1i (VICI).

Annotation predicted 14,246 protein-coding genes and 14,148 transcripts, with an average gene length of 2.5 kb. Reciprocal BLASTp (E-value threshold of 1 x 10^-5^) against the genome of *T. urticae* assigned gene descriptions to 67% of these genes. InterProScan and EggNOG analyses identified functional domains and assigned GO terms to 9,839 genes. Detailed read and assembly statistics are provided in Supplementary Table 1. The assembly and annotation are available under accession PRJNA1185618 and were used as the reference for the experiments described below.

### 3.2. Annotation of detoxification genes

Manual curation resulted in a comprehensive set of detoxification-related genes in *T. evansi*. The size and composition of these gene families were summarized and compared with those of *T. urticae* in Table 1.

**Table 1:**
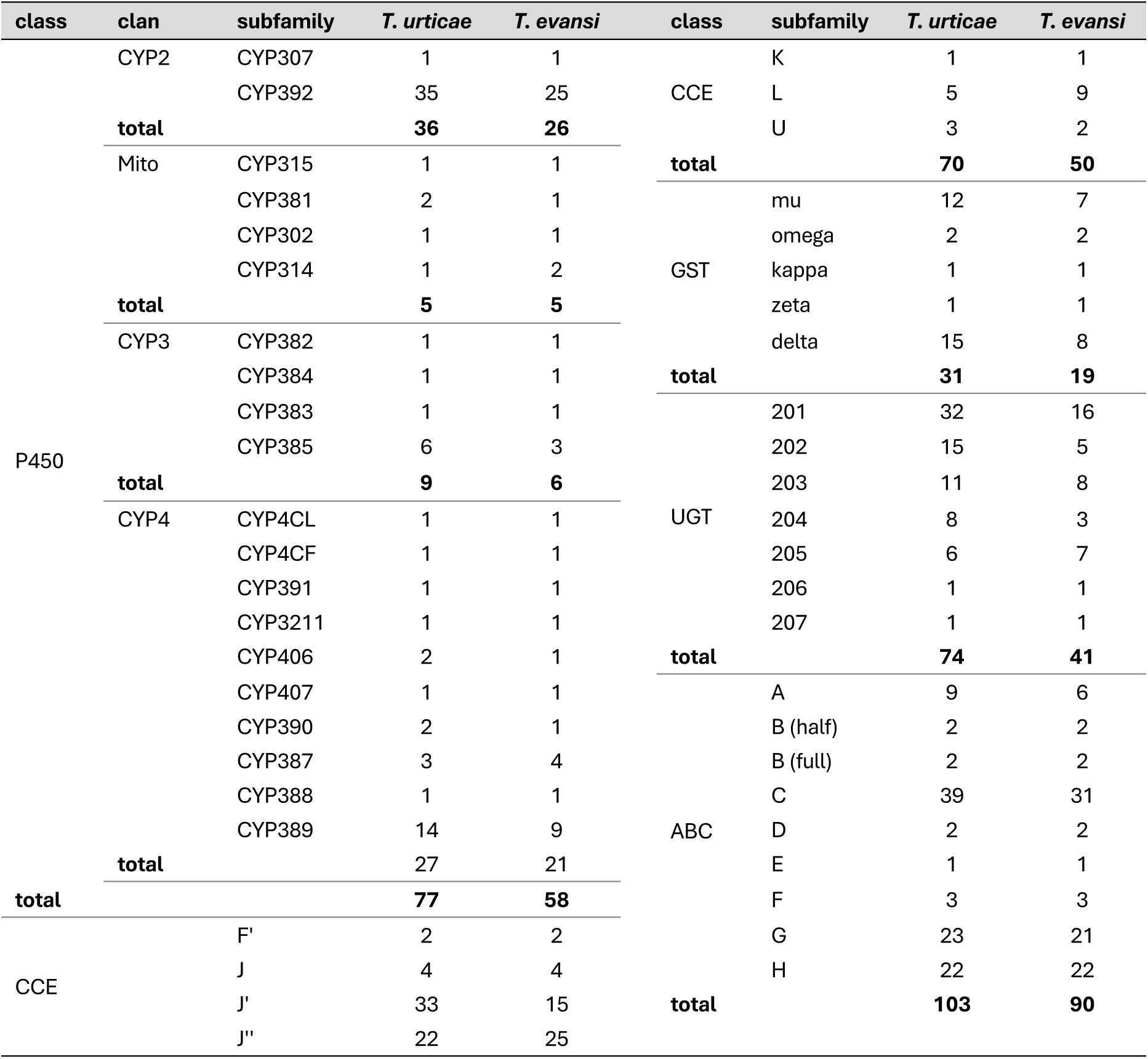
Comparative summary table of selected detoxification gene families in *T. urticae* and *T. evansi*. Pseudogenes were not included in the gene count.

The number of genes in all detoxification gene families was higher in *T. urticae* than in *T. evansi*. In particular, the UGT gene family was almost twice as large in *T. urticae* with 74 genes compared to 41 genes in *T. evansi*. This difference was mainly driven by expansions in *T. urticae* UGT201, UGT202, and UGT204 clades, while most other subfamilies were similar in size between species (Table 1). We identified 17 1:1 orthologues, 11 of which belonged to the UGT207, UGT206, UGT203, and UGT205 classes (Figure 2).

**Figure 2:**
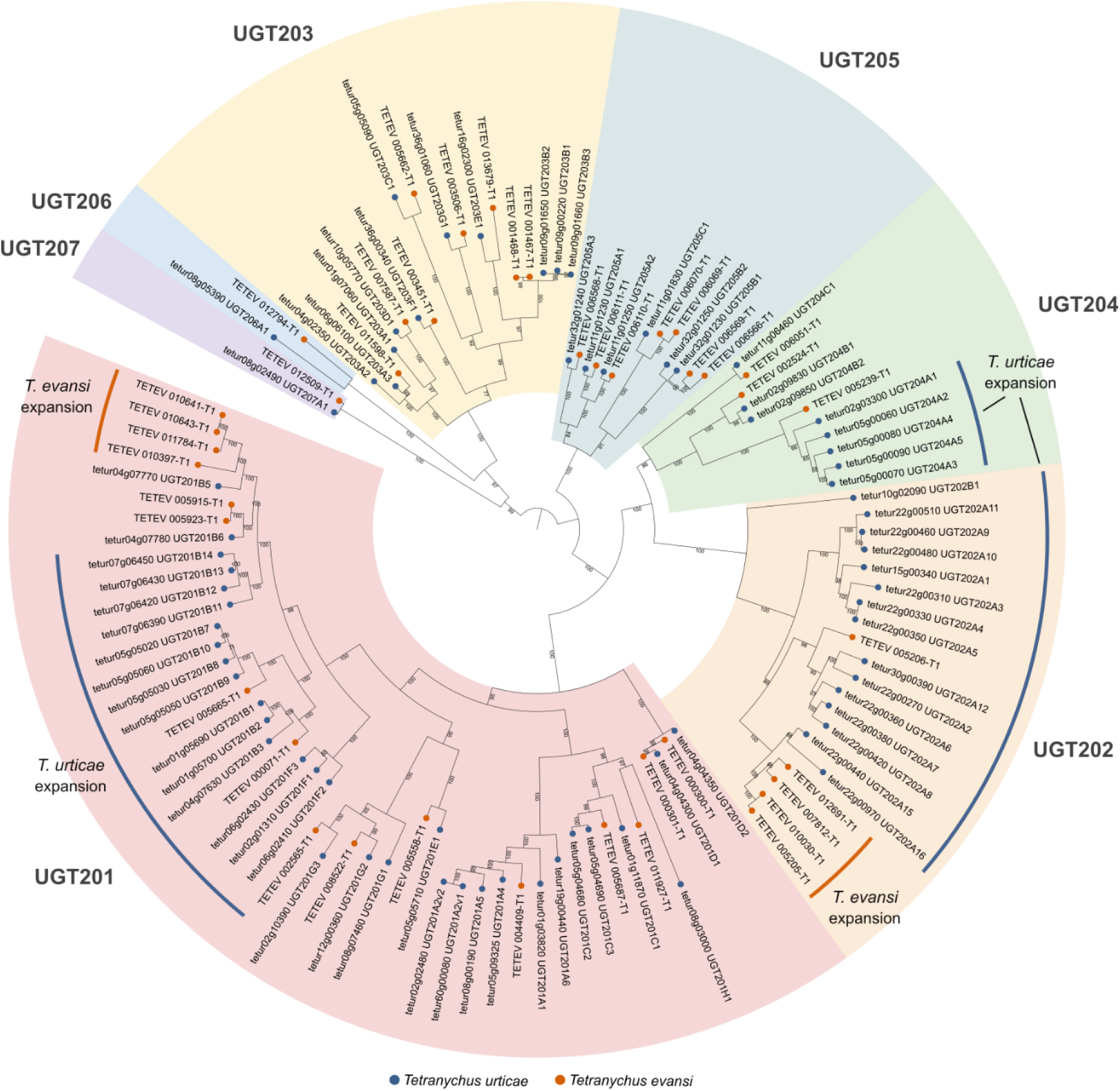
Phylogeny of UDP-glycosyltransferase (UGT) genes from *T. urticae* (blue) and *T. evansi* (red). Protein sequences were aligned using MAFFT (v7.526), and a maximum-likelihood tree was constructed with IQTREE (v3.0.1) using automatic model selection and 1,000 ultrafast bootstrap replicates. The resulting tree shows the UGTs grouped into the established *T. urticae* UGT clades. Species specific expansions are indicated.

*T. urticae* encoded 31 GST genes compared to 19 in *T. evansi*, largely due to expansions in the Mu and Delta classes (Supplementary Figure 2). A similar pattern was observed for CCE genes (85 vs 58) where the differences were mostly confined to the mite-specific J′ and J″ classes (Table 1, Supplementary Figure 3). In contrast, DOG gene numbers were more comparable between species (16 vs. 14) (Supplementary Figure 4). These genes, which are present in some mite species but absent from other animals, are involved in oxidative aromatic ring cleavage and are associated with detoxification and host plant adaptation (Dermauw et al., 2013a; Njiru et al., 2022). The ABC transporter gene family is only moderately larger in *T. urticae* (103 vs 90 in *T. evansi*), with the difference mainly attributable to the number of ABC-C genes, while little to no change was recorded in the number of genes in the other types of ABC transporter genes (Supplementary Figure 5). Similar to the other families, the P450s were less abundant in *T. evansi*, but with 25 1:1 orthologs (Table 1, Figure 3). The CYP392D and E subfamilies were expanded in *T. urticae*, while the CYP392A and B subfamilies appeared to have independently diversified in each species. The CYP314A1 gene encoding an ecdysone 20-hydroxylase was duplicated in *T. evansi*, an unusual occurrence in arthropods. Of note, we report two possible pseudogenes within the UGT gene family, namely *TETEV_013591* and *TETEV_006567*, while no pseudogenes were found in the other detox families.

**Figure 3:**
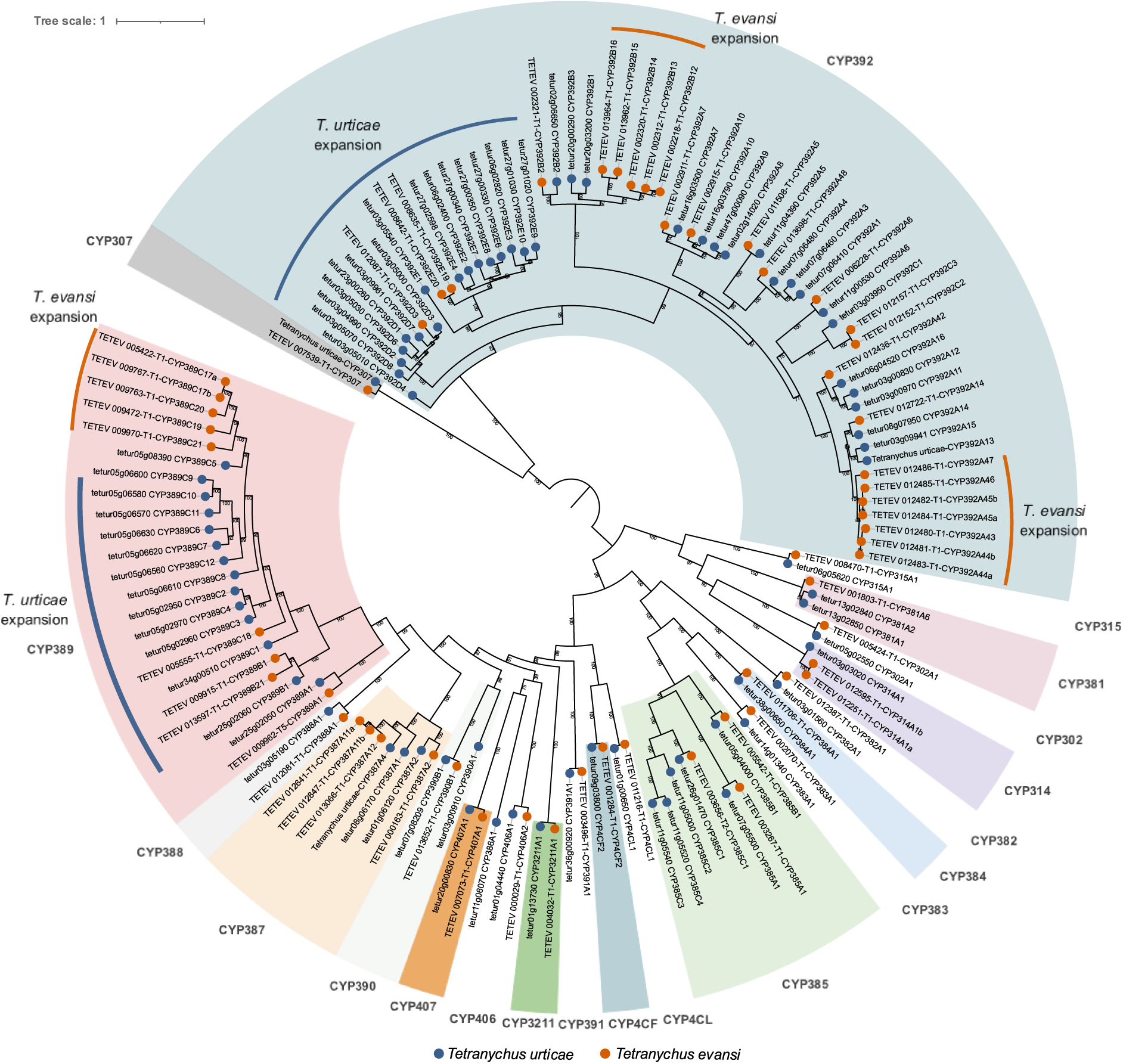
Phylogeny of cytochrome P450 mono-oxygenase genes from *T. urticae* (blue) and *T. evansi* (red). Protein sequences were aligned using MAFFT (v7.526), and a maximum-likelihood tree was constructed with IQTREE (v3.0.1) using automatic model selection and 1,000 ultrafast bootstrap replicates. The resulting tree shows the P450s grouped into the established clades and follow official nomenclature. Species-specific expansions are indicated.

### 3.3. Identification of orthologous carotenoid biosynthesis marker genes

The carotenoid biosynthesis genes of *T. evansi* are arranged in two conserved clusters on contig_1 (*TETEV_003851*, *TETEV_003852*) and contig_7 (*TETEV_011538*, *TETEV_011539*), with preserved gene order and orientation relative to *T. urticae* (Supplementary Figure 6) (Grbić et al., 2011a). A first tail-to-tail arrangement of the first cluster on chromosome 1 (*T. urticae*) and contig 1 (*T. evansi*) is consistent with fungi and aphids’ gene organization. The second cluster is a more complex arrangement of head-to-head *PD* genes followed by a reversed *CSC* gene in *T. urticae* (chromosome 3) while *T. evansi* possesses only a single *PD* and *CSC* gene on the contig 7 (Supplementary Figure 6). The two *CSC* genes have 1:1 orthologues in *T. urticae* and form a strongly supported monophyletic clade with aphid and fungal homologues (Figure 4), supporting their proposed origin via lateral gene transfer (Grbić et al., 2011a; Moran and Jarvik, 2010). In contrast, *PD* genes show a more complex evolutionary history: *TETEV_003852* is a 1:1 orthologue of *tetur01g11270*, whereas *TETEV_011538* clusters with two *T. urticae PD* genes and a pseudogene, indicating recent duplication and loss events. Based on this genomic context, the ortholog of *T. urticae tetur01g11270* (*TETEV_003852),* a gene whose disruption caused complete body discoloration in *T. urticae* (Bryon et al., 2017b; De Rouck et al., 2024) was selected as the target marker for CRISPR/Cas9-mediated gene editing (Figure 4).

**Figure 4:**
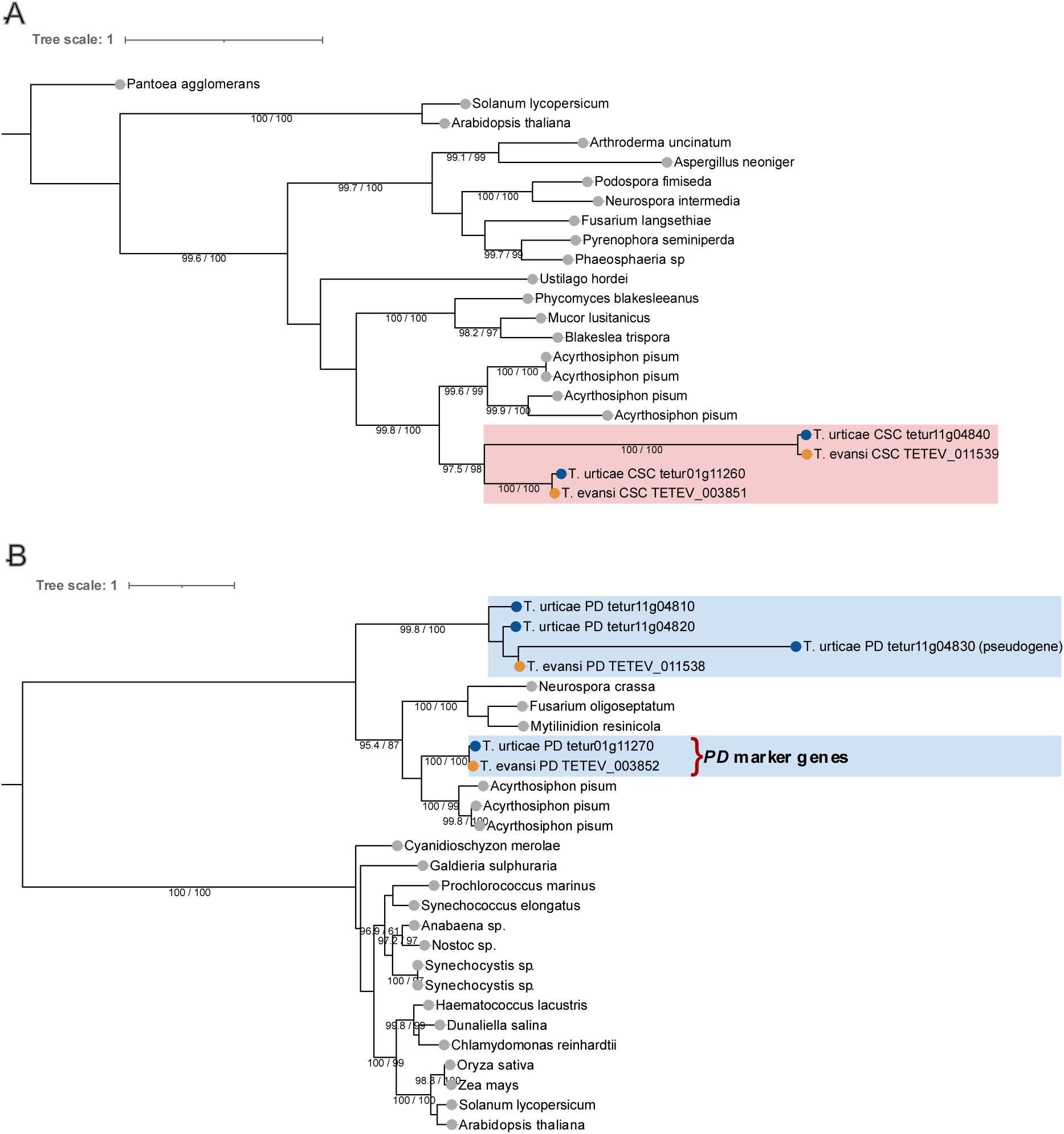
Maximum-likelihood phylogeny of the carotenoid cyclase/synthase (CSC) and phytoene desaturase (PD) proteins from selected green algae, land plants, fungi, and arthropod sequences. A) The CSC tree was rooted to the closest bacterial sequence of cyclases and synthases (*Pantoea*) (Grbić et al., 2011a). The two CSC proteins from *T. urticae* and *T. evansi* are 1:1 orthologues and form a monophyletic group with *Acyrthosiphon pisum* sequences. B) The tree was midpoint rooted. Bootstrap support (UFboot, 1,000 replicates) values are indicated at major nodes. Branch lengths are proportional to the number of amino acid substitutions per site.

### 3.4. Development of a gene-editing tool in *T. evansi*

Three independent CRISPR/Cas9 injections were performed using the SYNCAS formulation targeting the *PD* (*TETEV_003852*) gene in *T. evansi* (Figure 5). The resulting offspring were screened for mutant phenotypes associated with body discoloration, as previously described in *T. urticae* and other mite species (Bryon et al., 2017a; De Rouck et al., 2024). Sanger sequencing of G0 albino males confirmed the presence of CRISPR-induced mutations at the sgRNA target site. In addition, double peaks in the chromatograms of G0 females around the target region or downstream of the cut site indicated the presence of indel mutations.

**Figure 5.**
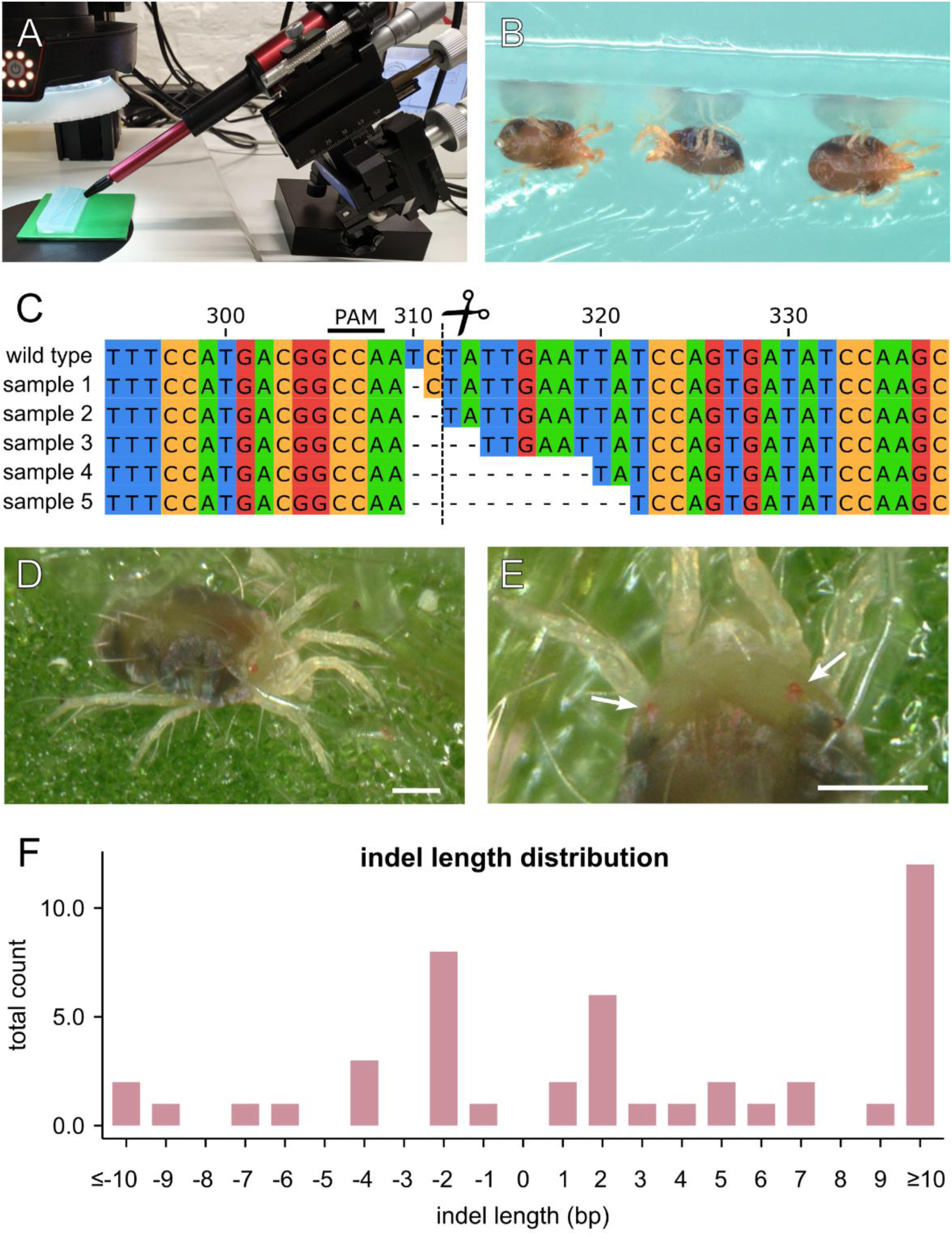
CRISPR-Cas9 setup and results targeting *phytoene desaturase* (*PD*). (A) Injection of set-up used to inject *T. evansi* mites (B) immobilized on agarose gel. C) Gene region in *PD* targeted by the sgRNA with indication of PAM- and cleavage-site. The alignment displays some of the mutations that were found in KO mutants. Numbers indicate nucleotide positions in the gene (including introns). (D-E) Adult G0 female showing partial body discoloration while retaining red eye pigmentation (arrows) due to maternal transfer of pigment by the WT mother. Scale bar represents 100 µm. (F) Frequency of insertion and deletion (indel) events detected in sequenced alleles. Negative values indicate deletions, positive values indicate insertions. Indel sizes are binned by base-pair length, with extreme categories grouped as ≤ −10 bp and ≥ 10 bp. The distribution shows a predominance of small indels, with 1–2 bp events being most frequent.

Analysis of indel length distribution at the *PD* target site revealed that small indels of 1–2 bp were predominant, accounting for roughly 17 events and representing most modification outcomes. Larger indels were comparatively rare, with most size classes above ±3 bp containing fewer than three events, except for insertions ≥10 bp, which comprised approximately 12 alleles, indicating a bias toward large insertions among rare editing outcomes (Figure 5F).

CRISPR/Cas9 editing of the *PD* gene yielded consistent mutation frequencies across the three independent replicates (Table 2). Among offspring collected 0-24 h post injection (TPI), overall editing rates ranged from 8.3% to 13.3%, while efficiency ranged from 7.3% to 15.9% in the 24-48 h interval. Mean editing efficiencies across replicates ranged from 9.6% to 14.7%. Mutations were detected in both sexes, although females generally exhibited higher editing frequencies than males. Together, the data demonstrates reliable and efficient CRISPR/Cas9-mediated gene editing in *T. evansi* using the SYNCAS formulation.

**Table 2:** CRISPR/Cas9 editing efficiency targeting the *PD* gene. . R, Replicate; TPI, Time Post Injection (h). “#offspring” indicates the total number of female (♀) and male (♂) offspring recovered for each TPI interval. “%mutant” represents the proportion of G0 individuals (♀/♂) carrying a CRISPR/Cas9-induced *PD* allele as determined by Sanger sequencing. “%mean editing” corresponds to the sex-independent editing efficiency over all offspring for the respective replicate.

| R | TPI (h) | #offspring<br>♀ | #offspring<br>♂ | #mutant<br>♀ | #mutant<br>♂ | Total | %mutant<br>♀ | %mutant<br>♂ | total<br>%editing | %mean<br>editing |
| --- | --- | --- | --- | --- | --- | --- | --- | --- | --- | --- |
| 1 | 0-24 | 113 | 128 | 18 | 14 | 32 | 15.93 | 10.94 | 13.3 | 9.6 |
|  | 24-48 | 203 | 193 | 9 | 20 | 29 | 4.43 | 10.36 | 7.3 |  |
| 2 | 0-24 | 95 | 15 | 10 | 4 | 14 | 10.53 | 26.67 | 12.7 | 12.9 |
|  | 24-48 | 106 | 16 | 15 | 1 | 16 | 14.15 | 6.25 | 13.1 |  |
| 3 | 0-24 | 5 | 7 | 0 | 1 | 1 | 0 | 14.29 | 8.3 | 14.7 |
|  | 24-48 | 49 | 14 | 7 | 3 | 10 | 14.29 | 21.43 | 15.9 |  |

Among females, one individual was found to be fully mutant, with both alleles carrying a KO allele at the CRISPR/Cas9 target site. This female was crossed with an albino G0 male to establish an albino line. Figure 6 shows the phenotype of each life stage for this *T. evansi* line. Notably, G0 mutants retained red eye pigmentation, whereas the remainder of their body lacked carotenoid-derived coloration (Figure 5D-E). This eye pigmentation was consistently lost in the subsequent generation, which suggests that G0 individuals retained carotenoid-like compounds from maternal origin.

**Figure 6:**
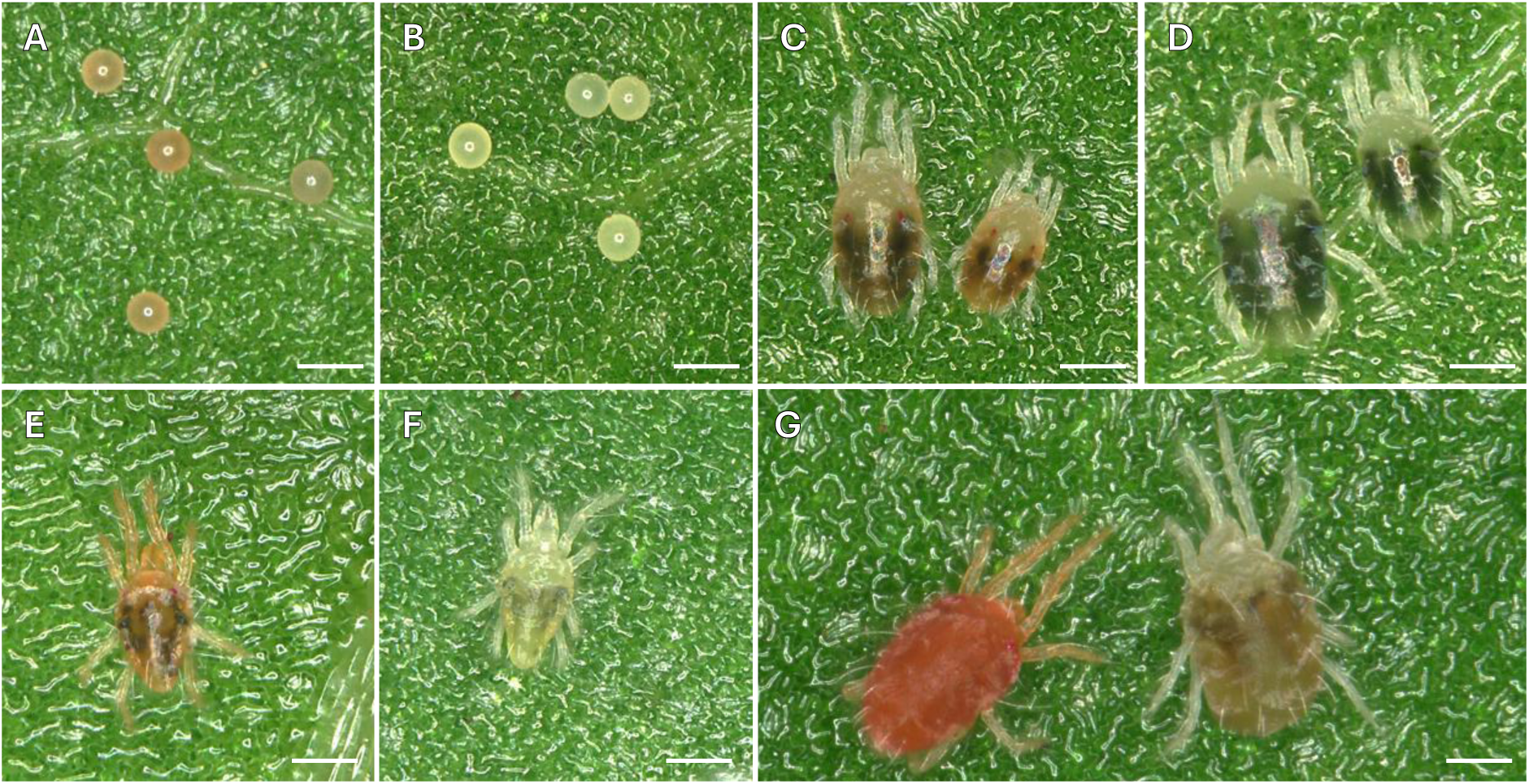
Phenotypes of wild-type *T. evansi* mites and mites with a *PD* knock-out. Shown are (A) eggs of the wild-type TEBA strain presenting a red pigmentation, (B) eggs of the TEBA *PD* KO strain presenting an albino phenotype characterized by a loss of red pigmentation, (C) and (D) are female (left) and male (right) of *T. evansi* mites at the teleiochrysalis stage from the wild-type and *PD* KO mites, respectively ; (E) wild-type adult male, (F) albino adult male, (G) wild-type adult female (left) and albino adult female (right). Scale bar represents 0.1 mm.

### 3.5. Introduction of the M918T substitution mutation in the VGSC gene

The M918T mutation was identified in the VGSC sequence of a *T. evansi* population exhibiting high levels of resistance to several pyrethroid insecticides and was hypothesized to be the causal resistance factor. To test the effect of this mutation on pyrethroid toxicity in *T. evansi*, we introduced this single nonsynonymous substitution into the susceptible TEBA strain using CRISPR/Cas9. The SYNCAS formulation was supplemented with an ssODN repair template carrying the A to C substitution, together with two silent marker mutations to avoid a second cleavage by Cas9 upon HDR (Supplementary Table 3).

A single experiment was performed in which 400 fertilized females were injected and allowed to lay eggs for 0–24 h and 24–48 h. For each oviposition time frame, 60–80 G0 couples were established and allowed to mate before being screened by Sanger sequencing to estimate CRISPR/Cas9 editing efficiency. In subsequent generations, genotyping was used to identify mutant individuals for directed crosses, ultimately resulting in four independent lines homozygous for the M918T mutation.

CRISPR/Cas9 editing of the *VGSC* gene resulted in mutations in both male and female G0 progeny across the two egg-laying windows. In offspring from eggs laid 0–24 h post-injection, 6.5% of males and 12.7% of females carried mutations. These mutations were characterized by primarily in-frame indels; in heterozygous females, the mutant allele was detected alongside the wild-type (WT) allele. Among the latter validated mutant individuals, 3.9% and 4.2% of males and females, respectively, carried the expected mutation with the silent markers, indicating HDR-mediated CRISPR/Cas9 edits (Table 3). In the 24–48 h cohort, mutation rates were 12.5% in males and 8.9% in females, with HDR detected in 8.9% of males and none of the females. Altogether, the mean KI frequency was 4.4%, indicating that SYNCAS-mediated CRISPR/Cas9 delivery can efficiently generate precise edits in *T. evansi*.

**Table 3:** CRISPR/Cas9 HDR efficiency for the modification of one nucleotide in the *VGSC* gene. TPI, Time Post Injection (h). “Total” represents the total number of individuals screened by Sanger sequencing among the offspring obtained from injected females for the corresponding TPI interval. “%mutants” represents the fraction of individuals for which the VGSC gene was edited and “%HDR” corresponds to the fraction of individuals presenting the expected nucleotide modification at the expected position (HDR).

| TPI (h) | Sex | Total | #mutants | #HDR | %mutants | %HDR | Total %mutants | Total %HDR | Mean %HDR |
| --- | --- | --- | --- | --- | --- | --- | --- | --- | --- |
| 0-24h | male | 77 | 5 | 3 | 6.5 | 3.9 | 9.5 | 4.1 | 4.4 |
|  | female | 71 | 9 | 3 | 12.7 | 4.2 |  |  |  |
| 24-48 | male | 56 | 7 | 5 | 12.5 | 8.9 | 10.9 | 5 |  |
|  | female | 45 | 4 | 0 | 8.9 | 0 |  |  |  |

Given the well-documented association of M918L in the VGSC with high levels of pyrethroid resistance in multiple arthropod species (De Rouck et al., 2023; Rinkevich et al., 2013), we also attempted to introduce this mutation into the *VGSC* gene of *T. evansi*. In contrast to the M918T experiment, editing M918L did not yield viable mutant males. G0 progeny consisted of both females and males, which were paired and allowed to mate. All G0 males were WT, whereas only a small number of G0 females carried the M918L mutation in heterozygous state. Male offspring from these crosses were consistently WT. Female offspring were isolated at the teleiochrysalis stage on bean leaf discs, allowed to oviposit, and subsequently genotyped. Although several G1 females carried the M918L mutation, none of their male progeny were mutant (Supplementary Figure 7B), preventing establishment of a mutant line. Collectively, these results suggest that the M918L mutation is either lethal or associated with a severe fitness cost in haploid males of *T. evansi*.

### 3.6. The contribution of M918T to pyrethroid resistance in *T. evansi*

Two of the four mutant lines established by gene-editing were used to assess the effect of the M918T mutation on pyrethroid toxicity in *T. evansi*. A full dose-response assay was performed on the WT TEBA strain to establish a LC_50_ for both type-I (bifenthrin) and type-II (beta-cyfluthrin) pyrethroids (Table 4). Bioassays with the VGSC^M918T-1^ and VGSC^M918T-2^ mutant lines showed a clear shift in LC_50_ with a value exceeding 2500 mg/L for bifenthrin, and a RR of 164 for VGSC^M918T-2^ towards beta-cyfluthrin.

**Table 4:** LC50 values of pyrethroid pesticides tested on WT and *VGSC^M918T^* TEBA lines.

| Compound | Strain | Slope ± se | LC<sub>50</sub> (95% CI) mg/L | χ<sup>2</sup> (df) | RR* |
| --- | --- | --- | --- | --- | --- |
| Bifenthrin | TEBA WT | 1.26 ± 0.11 | 2.26 (1.58 - 3.06) | 15.36 (26) | 1 |
|  | TEBA VGSC<sup>M918T-1</sup> |  | >2500 |  | >1,000 |
|  | TEBA VGSC<sup>M918T-2</sup> |  | >2500 |  | >1,100 |
| Beta-cyfluthrin | TEBA WT | 1.55 ± 0.13 | 0.96 (0.67 - 1.46) | 163.85 (18) | 1 |
|  | TEBA VGSC<sup>M918T-1</sup> | NA | NA | NA | NA |
|  | TEBA VGSC<sup>M918T-2</sup> | 2.66 ± 0.36 | 157.24 (60.24 - 307.85) | 31.59 (8) | 163,8 |

## 4. Discussion

Despite a long-standing status as invasive species and global threat to solanaceous crops, *T. evansi* lacked an annotated genome and genetic tools required for high-resolution functional investigations. In the present study, we present a draft genome assembly for *T. evansi* and establish efficient protocols for gene knockout and precision gene-editing, together laying the foundation for future genetic and genomic studies.

Using the Viçosa-1 line for seven generations of mother-son backcrossing, we created a line with near-complete homozygosity (Figure1). This lack of allelic complexity simplified genome assembly, yielding a highly contiguous reference (13 contigs, 94.8% BUSCO completeness). Although a genome assembly generated with Flye from a highly inbred line provides a strong long-read–based draft, especially for resolving repetitive regions and improving contiguity, assemblies can contain misjoins, collapsed repeats, or structural inaccuracies that are not easily detectable from contig statistics alone. Therefore, additional validation steps can confirm large-scale structural correctness and chromosomal organization. Incorporating Hi-C sequencing data can help anchor and orient contigs into chromosome-scale scaffolds while identifying potential misassemblies through interaction frequency patterns (Jia et al., 2019; Marcolungo et al., 2023). Similarly, optical mapping technologies (such as Bionano maps) offer an independent, long-range physical framework to validate structural consistency and detect inversions, translocations, or assembly gaps (Schwartz et al., 1993; Udall and Dawe, 2018). Last, and especially relevant for spider mites, allele frequency data from genetic mapping experiments can provide another complementary validation. This strategy was effectively demonstrated in Wybouw et al. (2019a) for *T. urticae*, where integration of allele frequency information available from bulk segregant analysis based genetic mapping experiments, helped identify assembly inconsistencies and refine genomic structure. Taken together, these validation approaches could substantially increase confidence in genome completeness and structural accuracy in the future.

We found an unexpected low fraction of heterozygous SNPs in the parental outbred Viçosa-1 strain, particularly when compared with its close relative, *T. urticae* (Cao et al., 2025; Grbić et al., 2011b; Xue et al., 2023). While *T. urticae* is known to maintain high levels of standing genetic variation even within long-term lab colonies (Wybouw et al., 2019a), *T. evansi* contains significantly less polymorphism. As Viçosa-1 represents the first *T. evansi* strain investigated for allelic variation, it is unknown whether this low polymorphism is an inherent characteristic of this lineage or a result of laboratory based population bottlenecks (Boubou et al., 2011; Habel and Schmitt, 2018). This observation is reinforced by the fact that our variant calling was performed using the same outbred population from which the inbred line for the genome assembly was derived, a process that inherently filters for lower allelic complexity. Sequencing DNA from more geographically and ecologically diverse populations of *T. evansi* will clarify this aspect in the future.

We carefully annotated over 280 genes across the major detoxification gene families including cytochrome P450 mono-oxygenases (P450), glutathione S-transferases (GST), carboxyl/choline esterases (CCE), ABC transporters as well as intradiol ring cleavage dioxygenases (DOG), the latter believed to have been acquired by mites from fungi via horizontal gene transfer (HGT) (Dermauw et al., 2013b; Njiru et al., 2022; Wybouw et al., 2014). While these families are classically linked to xenobiotic metabolism and host plant adaptation (Dermauw et al., 2020b; Després et al., 2007; Heckel, 2014), subfamily expansions are thought to dictate host plant breadth (Chen et al., 2023; Grbić et al., 2011b).

Comparing the specialist *T. evansi* to the extreme generalist *T. urticae* revealed significant genomic divergence. Differences between gene count in detox families might appear modest, but reflect changes in those subfamilies implicated in adaptation to toxins. This is most evident in the UGT201, UGT202 and UGT204 subfamilies, which have undergone massive expansions in *T. urticae*, with many enzymes directly linked to the degradation of plant secondary metabolites and acaricides (Snoeck et al., 2019). UGTs from these subfamilies exhibited broad flavonoid substrate specificity, including 7-hydroxyflavone, kaempferol and quercetin, as revealed by a high-throughput screening assay testing 44 compounds (Snoeck et al., 2019). Interestingly, two expansions also occurred in these subfamilies in *T. evansi* (UGT201 and UGT202), suggesting convergent functional importance. Furthermore, *T. evansi* evolved lineage-specific expansions of CYP genes (“CYP blooms”) in the CYP392 and CYP389 subfamilies, two clades that are largely expanded in *T. urticae* and are well-known for their roles in xenobiotic metabolism and metabolic resistance in both insects and mites (De Rouck et al., 2023; Nauen et al., 2022). Five members of the *T. urticae* CYP392 clade, namely CYP392A16, CYP392E10, CYP392A11, CYP392D2 and CYP392D8 demonstrated metabolic activity against more than 10 different pesticides (Demaeght et al., 2013; Fotoukkiaii et al., 2021; Lu et al., 2023; Riga et al., 2015, 2014; Tsakireli et al., 2024) while CYP389C16 was able to metabolize cyflumetofen and pyridaben (Feng et al., 2020). Although *T. urticae* possesses slightly more DOG genes than *T. evansi*, this difference is modest, and *T. evansi*-specific duplications of DOG genes were also observed. DOG enzymes function as a primary detoxification barrier in the mite gut. By catalyzing intradiol ring cleavage on catecholic aromatic rings, they effectively neutralize a broad spectrum of plant secondary metabolites. On tomato plants, which are particularly rich in these toxic phenolic compounds, DOG genes are essential for survival. Njiru et al. (2022) demonstrated that, upon transfer onto tomato plants, mites injected with *DOG11* or *DOG16* dsRNA exhibited significantly higher mortality compared to GFP dsRNA-injected controls. Therefore, DOG enzymes likely play a critical role in the specialization of *T. evansi* to solanaceous plants.

The availability of reverse genetic tools is crucial for functional studies of gene function and validation of the phenotypic impact of genetic variation. Here, we developed a genome-editing protocol based on the SYNCAS formulation, pioneered in *T. urticae* (De Rouck et al., 2024), which proved to be readily transferable to other spider mite species such as *T. evansi*. The phylogenetic proximity between *T. urticae* and *T. evansi* facilitated the prediction of phenotypic effects of gene knockout after *in silico* identification of orthologous marker genes such as the *phytoene desaturase* (*PD*), which allows easy visual screening of CRISPR events via eye and body pigmentation (Bryon et al., 2017a; De Rouck et al., 2024; Dermauw et al., 2020a). KO efficiency of the *PD* gene of up to 15.9% was obtained, which is of similar magnitude to that reported in other arthropods treated with the SYNCAS formulation, such as thrips (up to 30% KO efficiency in *Frankliniella occidentalis* <u>De Rouck et al., 2024</u>), whiteflies (20.0–39.1% in *Bemisia tabaci*, De Rouck et al., 2024; Mocchetti et al., 2025a), and 25% in *T. urticae* (De Rouck et al., 2024). However, during CRISPR pre-tests, we observed high mortality in *T. evansi* adult females when using the standard SYNCAS component doses originally developed for *T. urticae*. The toxicity of the formulation was decreased when the relative quantity of saponins was lowered from 2.6 to 1.3 μg/μL, which indicates that the toxic effect of saponins is more potent in *T. evansi*. This lower saponin concentration might explain the lower CRISPR efficiency. In addition, species-specific biological factors, such as differences in embryonic physiology, DNA repair dynamics, or tolerance to microinjection, as well as sgRNA- and target locus-dependent variability in editing efficiency, may also have contributed to the observed differences in CRISPR outcomes. Phenotypic differences between KO and WT mites were stage-consistent, though G0 mutants initially retained red eye pigmentation that vanished in subsequent generations. This transient phenotype likely reflects maternal contribution of pigment precursors or mRNA to the oocytes, a phenomenon similarly observed in *Plutella xylostella* and *Coccinella septempunctata* (Winters et al., 2014; Xu et al., 2020). The emergence of a full albino phenotype by the G1 generation confirms the depletion of these maternal reserves, thus validating that *PD* has a major role in eye coloration in *T. evansi* also, while at the same time demonstrating that the germ cells were edited allowing transmission of the mutation to the next generation. In contrast to *T. urticae*, where red eye pigmentation is not observed in the offspring of injected females, the persistence of red pigmentation in *T. evansi* G0 progeny likely reflects species-specific differences in pigment accumulation and/or maternal deposition. The markedly higher levels of pigment accumulation in *T. evansi* may delay the visible manifestation of complete albino phenotype until maternal stores are sufficiently depleted. Of particular note, we found one albino female among the G0 mutants across all replicated experiments, indicating that CRISPR events had occurred on both alleles. While this is consistent with previous results in *T. urticae* (De Rouck et al., 2024), it contrasts with results in predatory mites where, although editing efficiencies were lower than the ones reported here in *T. evansi,* multiple G0 homozygous females were identified.

CRISPR/Cas9 further offers a significant potential to unbiasedly validate resistance mutations in susceptible genetic backgrounds (İnak et al., 2024b; Mocchetti et al., 2026, 2025c). M918T in the VGSC is a target-site mutation known as *super-kdr* mutation, conferring high degrees of pyrethroid resistance (Dong et al., 2014; Field et al., 2017). In 2011, Nyoni et al. reported for the first time the presence of the M918T substitution in two Malawi populations of *T. evansi*, in the absence of any other *kdr* mutation. These populations showed 20 to 40-fold resistance to the pyrethroid bifenthrin compared to two susceptible field populations. Hence, as a first proof of principle for functional validation of a resistance mutation in this pest, M918T was selected to be introduced in a pyrethroid-susceptible *T. evansi* strain. We established two mutant lines from the TEBA strain, a *T. evansi* strain maintained on bean, a convenient host for performing crosses in CRISPR experiments and toxicity bioassays. HDR-based injections were performed only once and resulted in mean knock-in efficiency of 4.4%,. Phenotypic assessment of bifenthrin and beta-cyfluthrin toxicity of the TEBA *VGSC^M918T-1^* and *VGSC^M918T-2^* lines using dose-response assays resulted in LC_50_ values of >2,500 mg/L and 157.24 mg/mL respectively, which were >1,000- and 164-fold higher than the LC_50_ values obtained with the WT TEBA strain. The results obtained for bifenthrin shows considerable discrepancies of RR compared to the study of Nyoni et al. (2011), however this can be attributed to the higher basal level of tolerance to bifenthrin observed in that study. Our edited lines and the resistant field strains of Nyoni et al. show LC_50_ values of the same magnitude, substantiating the hypothesis that the M918T mutation alone can confer high levels of resistance to pyrethroid pesticides in *T. evansi*.

Interestingly, resistance conferred by M918T differ drastically between bifenthrin (RR >1000) and *beta*-cyfluthrin (RR = 164), likely indicating a different role of the mutation in affecting the binding of the two pyrethroids. The fact that specific amino acid substitutions can differentially affect the binding of different pyrethroids was already predicted through the VGSC model (O’Reilly et al., 2006). The residue at position 918, located at the bottom of the binding pocket where pyrethroids bind is known to be crucial in providing pyrethroid specificity. For instance, the methionine present at that position in arthropod VGSC is absent in mammals, and is considered one of the key differences causing the high selectivity of pyrethroids toward arthropods (Dong et al., 2014; Field et al., 2017; Zhorov and Dong, 2017). Docking studies investigating the effect of M918T in binding of different pyrethroids revealed that permethrin (Type I) and deltamethrin (Type II) would be highly affected, whereas binding of fenfluthrin (Type I) was not affected by the threonine at that position (Field et al., 2017; O’Reilly et al., 2006). Our data suggests that, in *T. evansi*, M918T alters more significantly the binding of bifenthrin than that of *beta*-cyfluthrin.

Both mite lines where the M918T mutation was fixed were easy to expand and no obvious fitness penalty was observed. Nevertheless, in the future it would be of interest to assess fitness costs. An unbiased approach that pairs well with the introgression of a resistance mutation via CRISPR/Cas9 is an experimental evolution setup, where, after an initial cross of TEBA (parental) and *VGSC^M918T^* lines (which share the same genetic background), WT and KI alleles can recombine and segregate freely within the resulting population in the absence of the selective pressure of the acaricide. Fitness cost assessment can then be done simply by monitoring allele frequency within the population over multiple generations (Mocchetti et al., 2025c; Njiru et al., 2023).

Attempts to establish a stable line carrying the alternative M918L substitution were unsuccessful, as no viable mutants were recovered among the haploid male offspring of viable heterozygous females. This finding suggests that M918L is hemizygous lethal in *T. evansi*. The M918L substitution has previously been reported in several other mite species, including *T. urticae*, *Dermanyssus gallinae*, *Varroa destructor*, *Phytoseiulus persimilis* and *Amblyseius swirskii* (Benavent-Albarracín et al., 2025; De Rouck et al., 2023). In these species, however, it was typically detected on the same haplotype as other *kdr* mutations, most commonly L925M/V, F1534L, and F1538I. Whether the presence of *kdr* mutations could mitigate the deleterious effects of M918L in *T. evansi* remains to be elucidated.

## 6. Credit authorship contribution statement

**Dries Amezian:** Data curation, Formal analysis, Validation, Investigation, Visualization, Methodology, Writing – original draft; **Femke De Graeve:** Data curation, Formal analysis, Validation, Investigation, Visualization, Methodology, Writing – original draft; **Rohith Mettumpurath Sasi:** Data curation, Formal analysis, Validation, Investigation, Visualization; **Lotte Vanhaecht:** Data curation, Formal analysis, Validation, Investigation, Visualization; **Antonio Mocchetti:** Investigation, Methodology, Writing – review and editing; **Ernesto Villacis-Perez:** Investigation, Methodology, Writing– review and editing; **Merijn Kant:** Funding acquisition; Writing – review and editing; **Sander De Rouck:** Validation, Investigation, Methodology, Writing – review and editing; **Thomas Van Leeuwen:** Conceptualization, Supervision, Funding acquisition, Writing – review and editing, Project administration.

## 7. Declaration of Generative AI and AI-assisted technologies in the writing process

During the preparation of this work the authors did not use generative AI or AI-assisted technologies in the writing process.

## Acknowledgements

Sander De Rouck is a post-doctoral fellow of FWO (grant 1237426N). This work was supported by Research Foundation Flanders (FWO) [1237426N, G0A1525N and G017923N], by a bilateral research cooperation between Flanders (FWO) – Vietnam (NAFOSTED) [grant G0E1221N], European Union’s Horizon 2020 research and innovation program (Grant agreement 101136611-NEXTGENBIOPEST), and the Dutch Research Council (NWO-VICI 19391).

## 8. Data Availibility

The genome assembly and annotation, along with the raw ONT sequencing data, have been deposited in NCBI under BioProject accession number PRJNA1185618. The whole-genome sequencing (WGS) data used for variant calling and RNA sequencing data used for annotation are available under BioProject accession numbers [*in process*] and [*in process*], respectively.

## Supplementary material

### Supplementary Tables

**Supplementary Table 1:** Summary of sequencing input, genome assembly metrics, and gene annotation statistics for the Vicosa-1 inbred genome assembly.

| ONT reads (used for assembly; post-filtering (see Material and Methods)) |  |
| --- | --- |
| Total Gbases | 3.4 |
| Total number of reads |  |
| N50 | 19,227 |
| N90 | 11,155 |
| Genome assembly |  |
| Total size (bp) | 89,329,059 |
| Mean coverage | 36 |
| Number of fragments | 13 |
| Fragments N50 (bp) | 18,641,227 |
| Fragments N90 (bp) | 5,924,920 |
| Largest fragment (bp) | 31,544,786 |
| Number of scaffolds | 0 |
| Number of contigs | 13 |
| GC content (%) | 31.89 |
| L50 | 2 |
| L90 | 6 |
| Repetitive elements (%) | NA |
| Annotation |  |
| Number of genes | 14,148 |
| Average gene length | 2,582 |
| Average CDS length | 1,299 |
| Average exon length | 507 |
| Number of single exon gene | 2,501 |
| Genes with GO term assigned | 8,819 |
| Genes with IPR domain assigned | 10,091 |
| Genes with PFAM domain assigned | 8,630 |

**Supplementary Table 2:** list of phytoene desaturase and carotenoid cyclase/synthase genes and IDs from diverse clades used for construction of the maximum-likelihood phylogenetic tree.

| Gene | Species | GenBank accession | Length (aa*) |
| --- | --- | --- | --- |
| Phytoene desaturase | Arabidopsis thaliana | NP_193157.1 | 566 |
|  | Zea mays | NP_001338939.1 | 571 |
|  | Solanum lycopersicum | CAB59726.1 | 583 |
|  | Oryza sativa | NP_001404184.1 | 584 |
|  |  | XP_029343890.1 | 526 |
|  | Acyrtosiphon pisum | XP_001943225.2 | 528 |
|  |  | XP_001950764.1v | 528 |
|  | Lachnellula hyalina | XP_031009100.1 | 596 |
|  | Mytilinidion resinicola | XP_033574400.1 | 885 |
|  | Chlamydomonas reinhardtii | AAT38476.1 | 564 |
|  | Haematococcus lacustris | AKO71906.1 | 569 |
|  | Synechocystis sp. PCC 6803 | CAA44452.1 | 472 |
|  | Dunaliella salina | ADD52599.1v | 582 |
|  | Neurospora crassa OR74A | XP_964713.1 | 595 |
|  | Fusarium oligoseptatum | RSM12532.1 | 870 |
|  | Synechocystis sp. PCC 6803 | BAA17847.1 | 472 |
|  | Synechococcus elongatus PCC 7942 | CAA39004.1 | 474 |
|  | Nostoc sp. PCC 7120 | RUR82889.1 | 479 |
|  | Anabaena sp. AL93 | OBQ17212.1 | 479 |
|  | Prochlorococcus marinus subsp. pastoris str. CCMP1986 | CAE18603.1 | 473 |
|  | Cyanidioschyzon merolae strain 10D | BAM80526.1 | 575 |
|  | Galdieria sulphuraria | XP_005704373.1 | 539 |
| Carotenoid synthase/cyclase |  | XP_003241668 | 599 |
|  | Acyrtosiphon pisum | XP_001950787 | 588 |
|  |  | NP_001171302 | 531 |
|  | Blakeslea trispora | AAO46895.1 | 608 |
|  | Ustilago hordei | CCF48514.1 | 701 |
|  | Phycomyces blakesleeanus | KAL0093072.1 | 602 |
|  | Aspergillus niger | PYH30037.1 | 589 |
|  | Arthroderma uncinatum | XP_033405613.1 | 428 |
|  | Pantoea agglomerans | AAA64982.1 | 309 |
|  | Mucor lusitanicus | KAL0143141.1 | 614 |
|  | Neurospora intermedia | KAL0475314.1 | 602 |
|  | Podospora fimseda | KAK4229975.1 | 586 |
|  | Phaeosphaeria sp. | KAH7405766.1 | 585 |
|  | Pyrenophora seminiperda | RMZ72926.1 | 861 |
|  | Fusarium langsethiae | KPA39208.1 | 581 |
|  | Solanum lycopersicum | NP_001334767.1 | 412 |
|  | Arabidopsis thaliana | NP_197225.1 | 422 |

\* aa: amino acid

**Supplementary Table 3:**
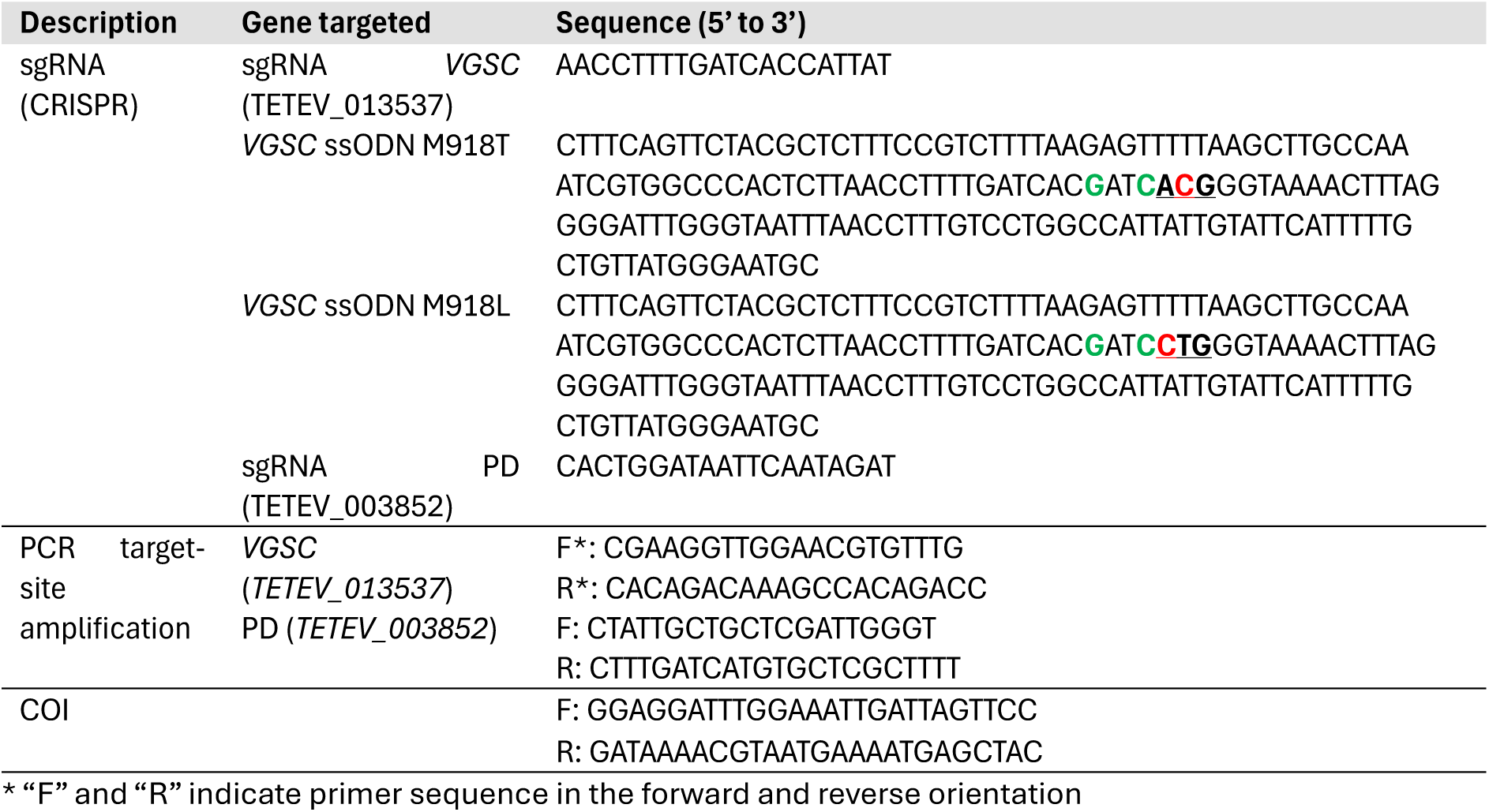
list of oligos and primers.

### Supplementary Figures

**Supplementary Figure 1:**
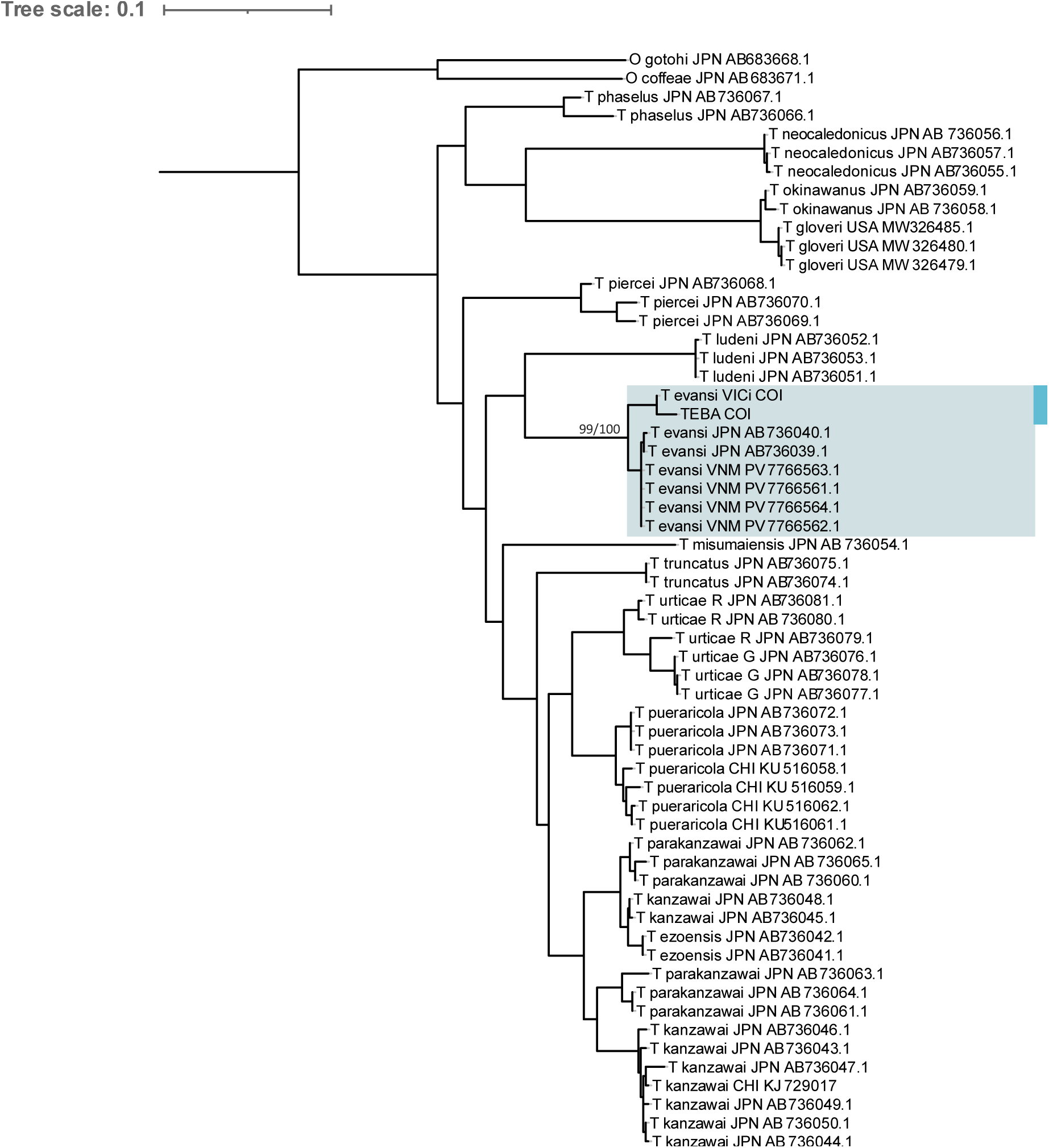
Maximum-likelihood phylogenetic tree of spider mite populations based on *COI* fragments. As the geographical origin from the *T. evansi* bean-adapted (TEBA) strain was not recorded, its *COI* marker was amplified and aligned to *COI* sequences from *T. evansi* Viçosa-1i (VICi) and various spider mite populations (Mocchetti et al., 2025b) using MAFFT (v7.525). IQTREE (v3.0.1) was used to build a phylogenetic tree from this alignment, which confirms that TEBA is a population of *T. evansi* spider mites.

**Supplementary Figure 2:**
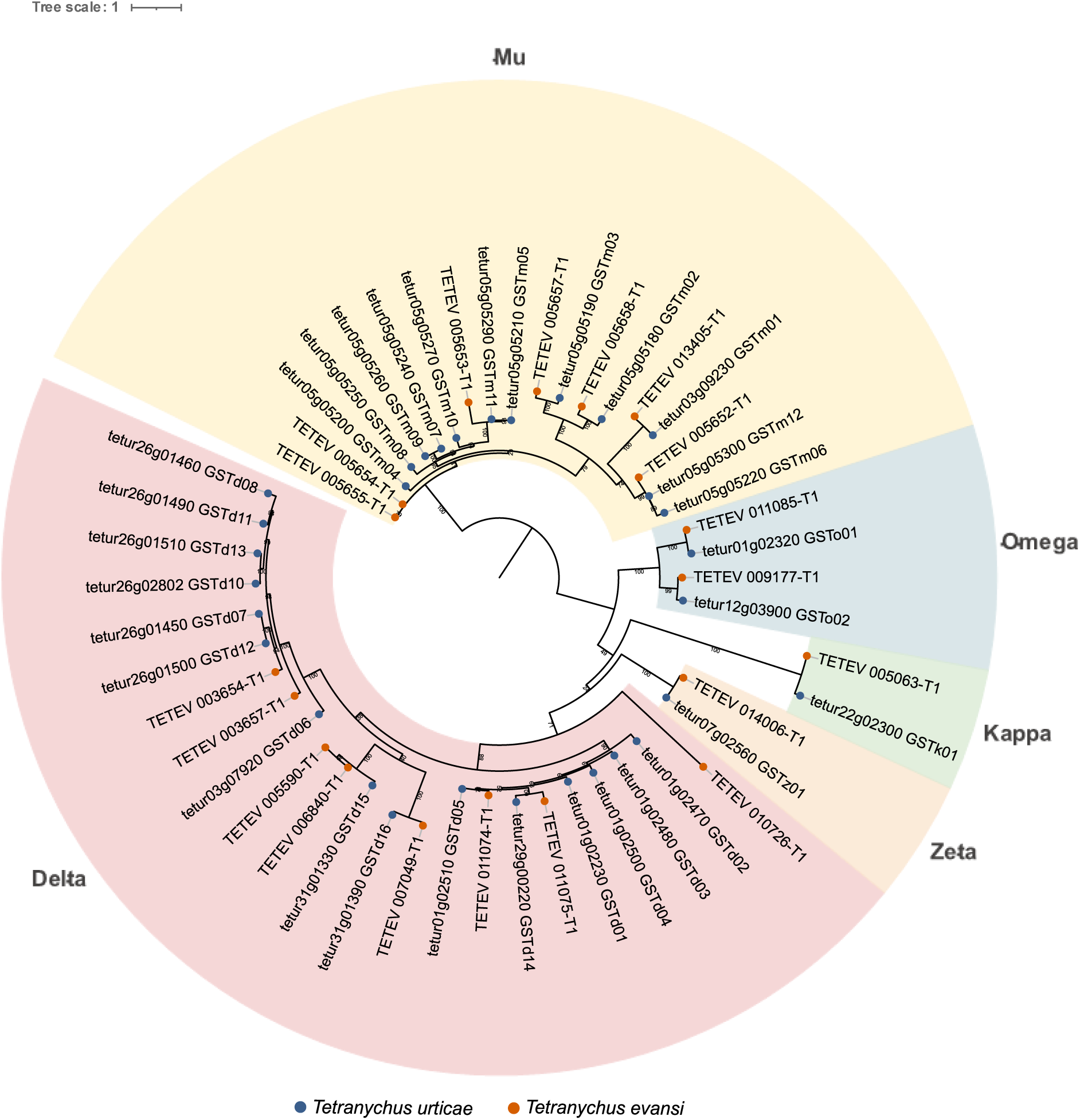
Phylogeny of glutathione-S-transferase (GST) genes from *T. urticae* (blue) and *T. evansi* (red). Protein sequences were aligned using MAFFT (v7.526), and a maximum-likelihood tree was constructed with IQTREE (v3.0.1) using automatic model selection and 1,000 ultrafast bootstrap replicates. The resulting tree shows the GSTs grouped into the established *T. urticae* GST clades.

**Supplementary Figure 3:**
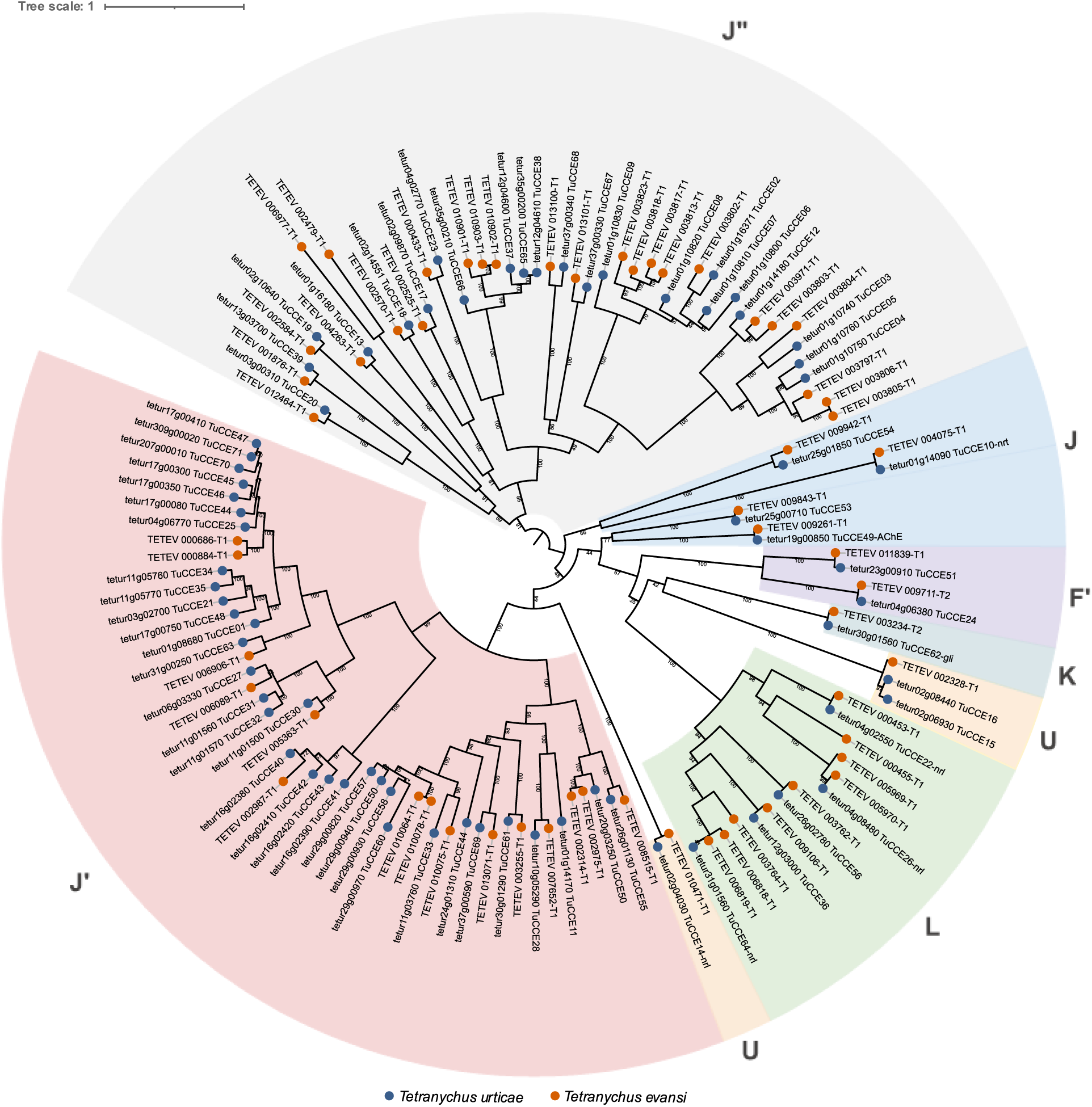
Phylogeny of carboxyl/choline esterase (CCEs) genes from *T. urticae* (blue) and *T. evansi* (red). Protein sequences were aligned using MAFFT (v7.526), and a maximum-likelihood tree was constructed with IQTREE (v3.0.1) using automatic model selection and 1,000 ultrafast bootstrap replicates. The resulting tree shows the CCEs grouped into the established *T. urticae* CCE clades.

**Supplementary Figure 4:**
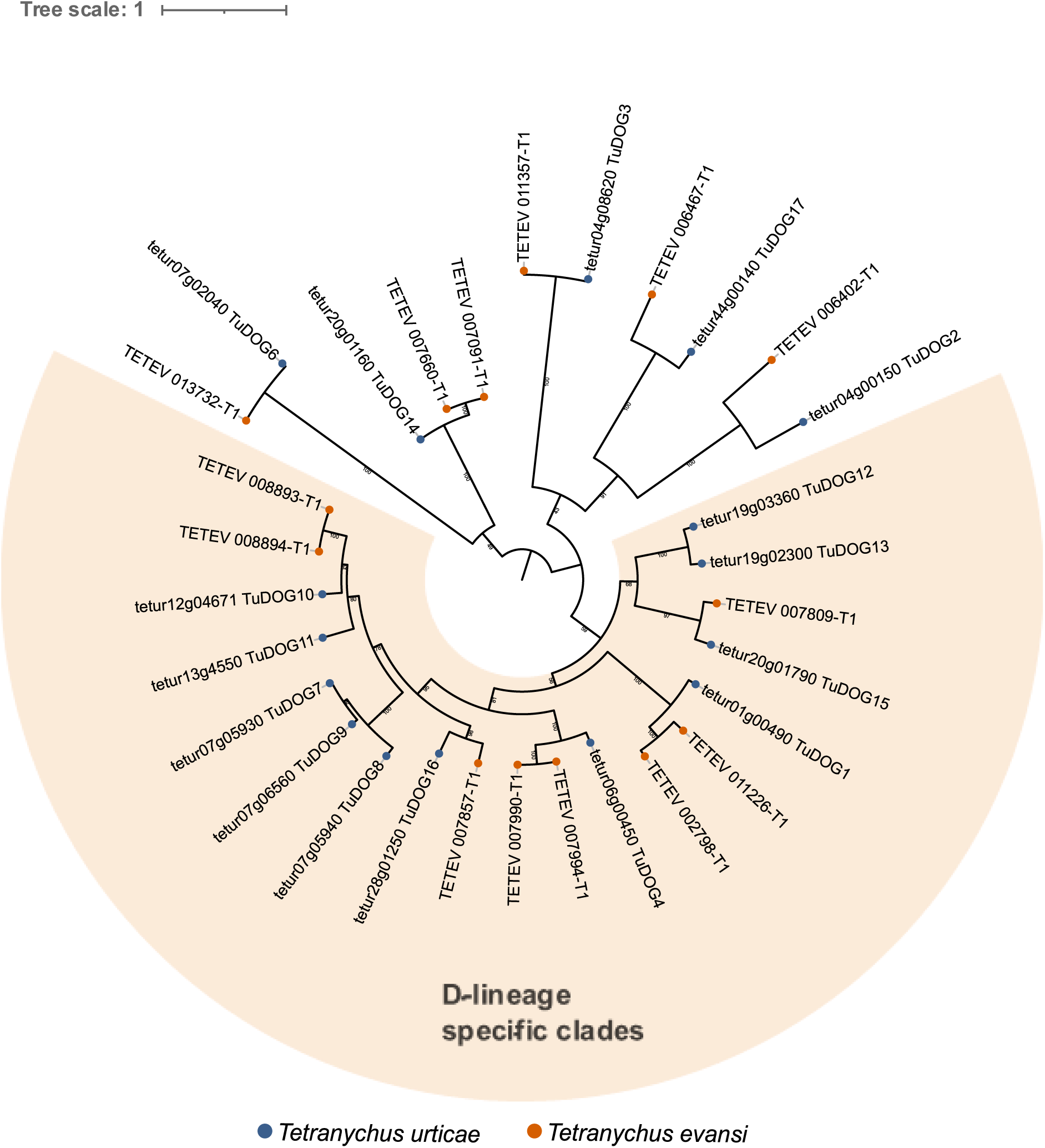
Phylogeny of Intradiol ring cleavage dioxygenases (DOG) genes from *T. urticae* (blue) and *T. evansi* (red). Protein sequences were aligned using MAFFT (v7.526), and a maximum-likelihood tree was constructed with IQTREE (v3.0.1) using automatic model selection and 1,000 ultrafast bootstrap replicates.

**Supplementary Figure 5:**
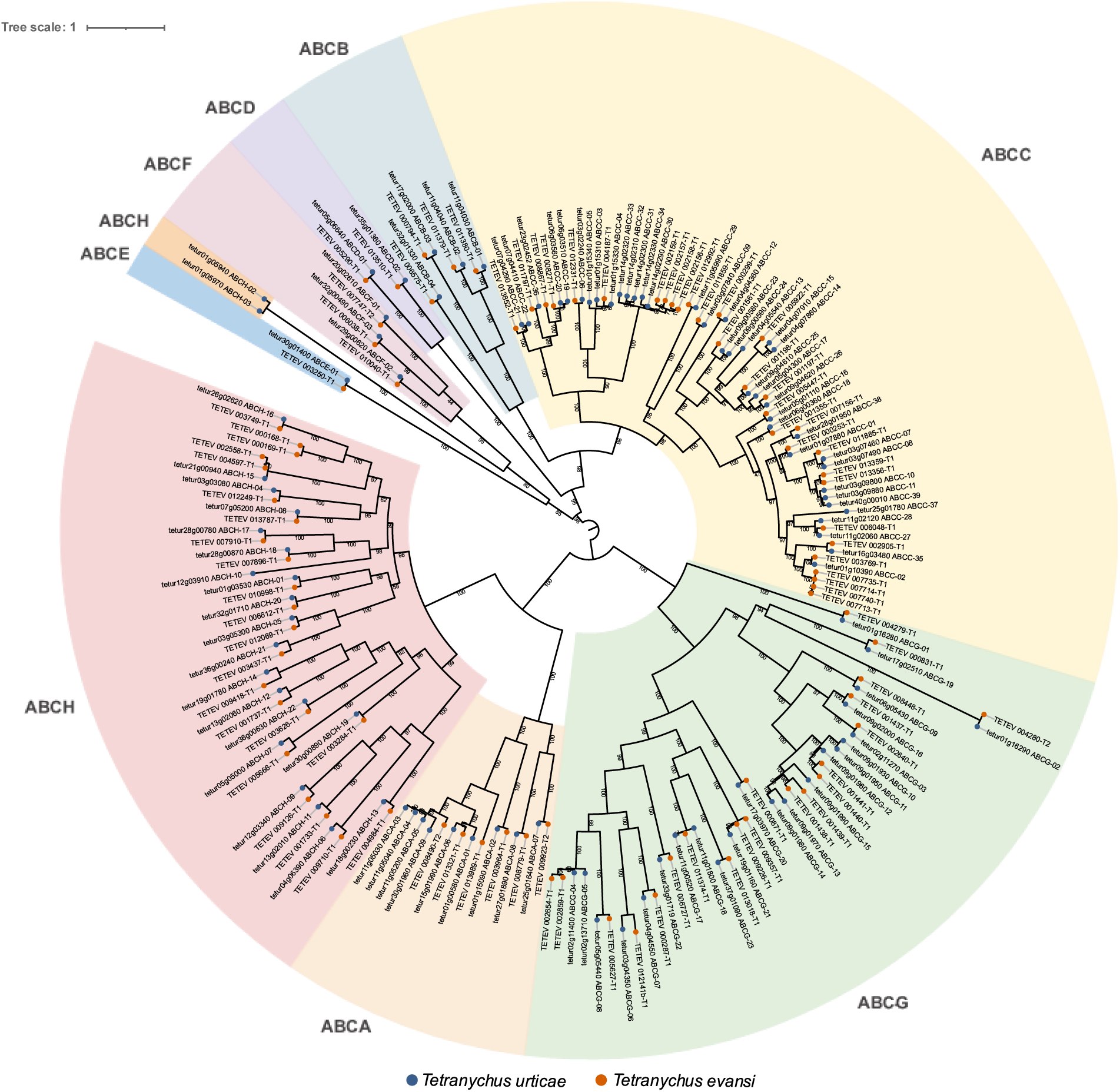
Phylogeny of ABC-transporter (ABC) genes from *T. urticae* (blue) and *T. evansi* (red). Protein sequences were aligned using MAFFT (v7.526), and a maximum-likelihood tree was constructed with IQTREE (v3.0.1) using automatic model selection and 1,000 ultrafast bootstrap replicates. The resulting tree shows the ABCs grouped into the established *T. urticae* ABC clades.

**Supplementary Figure 6:**
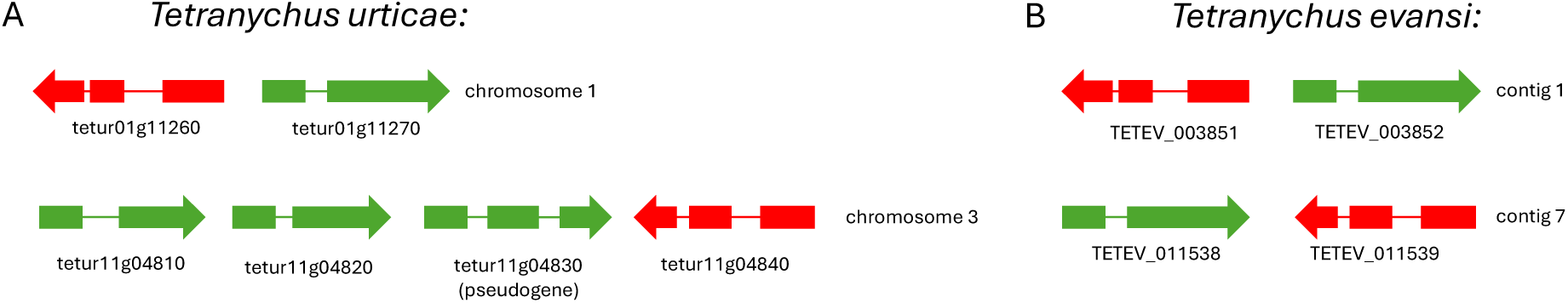
Organization of carotenoid biosynthesis gene clusters in *T. urticae* and *T. evansi*.

**Supplementary Figure 7:**
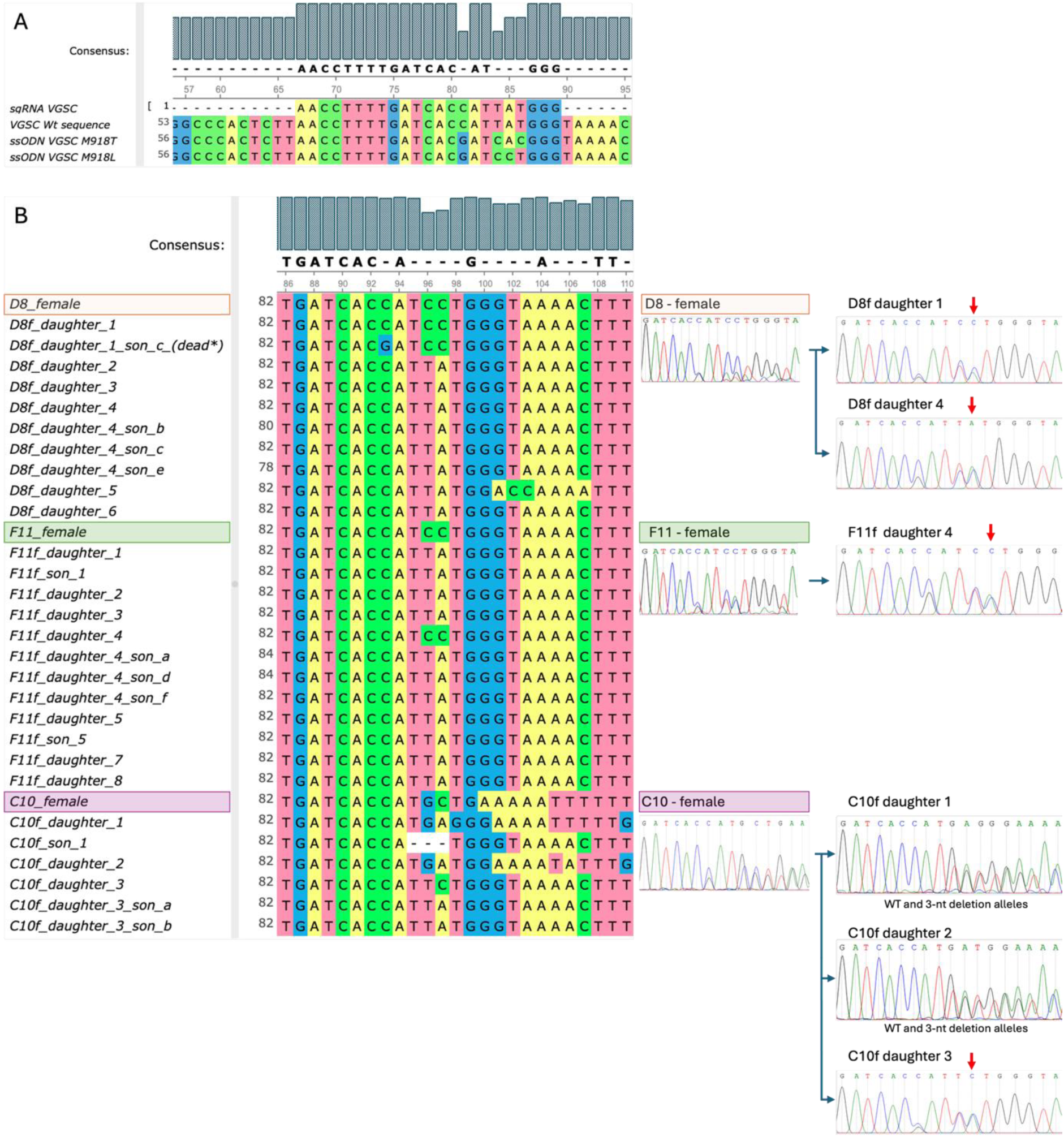
Genetic screening of M918L gene-edited mites. A) Alignment of the *VGSC* genomic sequence at the CRISPR/Cas9 target site (WT), the sgRNA, and the ssODN templates designed to introduce the M918T and M918L substitutions. B) Sanger sequencing-based screening of G0 and subsequent generations in the M918L experiment. The alignment shows sequences from selected females and males. Three G0 females carrying the heterozygous M918L mutation are highlighted in orange (D8_female), green (F11_female), and purple (C10_female), with corresponding chromatograms shown to the right. Female G1 progeny derived from these individuals were sequenced (e.g. F11f_daughter_1, F11f_daughter_2, etc.) after oviposition for a few days. Although some G1 females carried the M918L mutation (e.g. F11f_daughter_4), none of their male offspring were mutant (see alignment). Red arrows above chromatograms indicate the position of the A→C substitution corresponding to the M918L mutation.

## References

Alavijeh, E.S., Khajehali, J., Snoeck, S., Panteleri, R., Ghadamyari, M., Jonckheere, W., Bajda, S., Saalwaechter, C., Geibel, S., Douris, V., Vontas, J., Van Leeuwen, T., Dermauw, W., 2020. Molecular and genetic analysis of resistance to METI-I acaricides in Iranian populations of the citrus red mite Panonychus citri. Pesticide Biochemistry and Physiology 164, 73–84. 10.1016/j.pestbp.2019.12.009

Azandémè-Hounmalon, G.Y., Affognon, H.D., Komlan, F.A., Tamò, M., Fiaboe, K.K.M., Kreiter, S., Martin, T., 2015. Farmers’ control practices against the invasive red spider mite, *Tetranychus evansi* Baker & Pritchard in Benin. Crop Protection 76, 53–58. 10.1016/j.cropro.2015.06.007

Azandeme-Hounmalon, G.Y., Sikirou, R., Onzo, A., Fiaboe, K.K.M., Tamo, M., Kreiter, S., Martin, T., 2022. Re-assessing the pest status of *Tetranychus evansi* (Acari: Tetranychidae) on solanaceous crops and farmers control practices in Benin. Journal of Agriculture and Food Research 10, 100401. 10.1016/j.jafr.2022.100401

Bajda, S., Dermauw, W., Panteleri, R., Sugimoto, N., Douris, V., Tirry, L., Osakabe, M., Vontas, J., Van Leeuwen, T., 2017. A mutation in the PSST homologue of complex I (NADH:ubiquinone oxidoreductase) from *Tetranychus urticae* is associated with resistance to METI acaricides. Insect Biochemistry and Molecular Biology 80, 79–90. 10.1016/j.ibmb.2016.11.010

Benavent-Albarracín, L., Pérez-Hedo, M., Alonso-Valiente, M., Catalán, J., Urbaneja, A., González-Cabrera, J., 2025. Response of Amblyseius swirskii to deltamethrin. Pest Management Science 81, 2800–2811. 10.1002/ps.8647

Boubou, A., Migeon, A., Roderick, G.K., Navajas, M., 2011. Recent emergence and worldwide spread of the red tomato spider mite, Tetranychus evansi: genetic variation and multiple cryptic invasions. Biol Invasions 13, 81–92. 10.1007/s10530-010-9791-y

Bryon, A., Kurlovs, A.H., Dermauw, W., Greenhalgh, R., Riga, M., Grbić, M., Tirry, L., Osakabe, M., Vontas, J., Clark, R.M., Van Leeuwen, T., 2017a. Disruption of a horizontally transferred phytoene desaturase abolishes carotenoid accumulation and diapause in Tetranychus urticae. Proceedings of the National Academy of Sciences 114, E5871–E5880. 10.1073/pnas.1706865114

Bryon, A., Kurlovs, A.H., Dermauw, W., Greenhalgh, R., Riga, M., Grbić, M., Tirry, L., Osakabe, M., Vontas, J., Clark, R.M., Van Leeuwen, T., 2017b. Disruption of a horizontally transferred phytoene desaturase abolishes carotenoid accumulation and diapause in Tetranychus urticae. Proceedings of the National Academy of Sciences 114, E5871–E5880. 10.1073/pnas.1706865114

Camacho, C., Coulouris, G., Avagyan, V., Ma, N., Papadopoulos, J., Bealer, K., Madden, T.L., 2009. BLAST+: architecture and applications. BMC Bioinformatics 10, 421. 10.1186/1471-2105-10-421

Cao, L.-J., Chen, J.-C., Thia, J.A., Schmidt, T.L., Ffrench-Constant, R., Zhang, L.-X., Yang, Y., Yuan, M.-C., Zhang, J.-Y., Zhang, X.-Y., Yang, Q., Gong, Y.-J., Li, H., Chen, X., Hoffmann, A.A., Wei, S.-J., 2025. Recurrent mutations drive the rapid evolution of pesticide resistance in the two-spotted spider mite Tetranychus urticae. eLife 14, RP106288. 10.7554/eLife.106288

Carvalho, R., Yang, Y., Field, L.M., Gorman, K., Moores, G., Williamson, M.S., Bass, C., 2012. Chlorpyrifos resistance is associated with mutation and amplification of the acetylcholinesterase-1 gene in the tomato red spider mite, *Tetranychus evansi*. Pesticide Biochemistry and Physiology, Special Issue: Molecular Approaches to Pest Control, Toxicology and Resistance 104, 143–149. 10.1016/j.pestbp.2012.05.009

Charamis, J., Dermauw, W., Van Leeuwen, T., Vontas, J., Feyereisen, R., 2025. The arthropod P450 Enchiridion: An integrated web resource for research on P450s. Insect Biochemistry and Molecular Biology 183, 104377. 10.1016/j.ibmb.2025.104377

Chen, L., Yu, X.-Y., Xue, X.-F., Zhang, F., Guo, L.-X., Zhang, H.-M., Hoffmann, A.A., Hong, X.-Y., Sun, J.-T., 2023. The genome sequence of a spider mite, Tetranychus truncatus, provides insights into interspecific host range variation and the genetic basis of adaptation to a low-quality host plant. Insect Science 30, 1208–1228. 10.1111/1744-7917.13212

Concordet, J.-P., Haeussler, M., 2018. CRISPOR: intuitive guide selection for CRISPR/Cas9 genome editing experiments and screens. Nucleic Acids Res 46, W242–W245. 10.1093/nar/gky354

De Rouck, S., İnak, E., Dermauw, W., Van Leeuwen, T., 2023. A review of the molecular mechanisms of acaricide resistance in mites and ticks. Insect Biochemistry and Molecular Biology 159, 103981. 10.1016/j.ibmb.2023.103981

De Rouck, S., Mocchetti, A., Dermauw, W., Van Leeuwen, T., 2024. SYNCAS: Efficient CRISPR/Cas9 gene-editing in difficult to transform arthropods. Insect Biochemistry and Molecular Biology 165, 104068. 10.1016/j.ibmb.2023.104068

Demaeght, P., Dermauw, W., Tsakireli, D., Khajehali, J., Nauen, R., Tirry, L., Vontas, J., Lümmen, P., Van Leeuwen, T., 2013. Molecular analysis of resistance to acaricidal spirocyclic tetronic acids in Tetranychus urticae: CYP392E10 metabolizes spirodiclofen, but not its corresponding enol. Insect Biochemistry and Molecular Biology 43, 544–554. 10.1016/j.ibmb.2013.03.007

Dermauw, W., Jonckheere, W., Riga, M., Livadaras, I., Vontas, J., Van Leeuwen, T., 2020a. Targeted mutagenesis using CRISPR-Cas9 in the chelicerate herbivore *Tetranychus urticae*. Insect Biochemistry and Molecular Biology 120, 103347. 10.1016/j.ibmb.2020.103347

Dermauw, W., Van Leeuwen, T., Feyereisen, R., 2020b. Diversity and evolution of the P450 family in arthropods. Insect Biochemistry and Molecular Biology 127, 103490. 10.1016/j.ibmb.2020.103490

Dermauw, W., Wybouw, N., Rombauts, S., Menten, B., Vontas, J., Grbić, M., Clark, R.M., Feyereisen, R., Van Leeuwen, T., 2013a. A link between host plant adaptation and pesticide resistance in the polyphagous spider mite Tetranychus urticae. Proceedings of the National Academy of Sciences 110, E113–E122. 10.1073/pnas.1213214110

Dermauw, W., Wybouw, N., Rombauts, S., Menten, B., Vontas, J., Grbić, M., Clark, R.M., Feyereisen, R., Van Leeuwen, T., 2013b. A link between host plant adaptation and pesticide resistance in the polyphagous spider mite Tetranychus urticae. Proceedings of the National Academy of Sciences 110, E113–E122. 10.1073/pnas.1213214110

Després, L., David, J.-P., Gallet, C., 2007. The evolutionary ecology of insect resistance to plant chemicals. Trends in Ecology & Evolution 22, 298–307. 10.1016/j.tree.2007.02.010

Ding, T.-B., Zhong, R., Jiang, X.-Z., Liao, C.-Y., Xia, W.-K., Liu, B., Dou, W., Wang, J.-J., 2015. Molecular characterisation of a sodium channel gene and identification of a Phe1538 to Ile mutation in citrus red mite, Panonychus citri. Pest Management Science 71, 266–277. 10.1002/ps.3802

Dong, K., Du, Y., Rinkevich, F., Nomura, Y., Xu, P., Wang, L., Silver, K., Zhorov, B.S., 2014. Molecular biology of insect sodium channels and pyrethroid resistance. Insect Biochemistry and Molecular Biology 50, 1–17. 10.1016/j.ibmb.2014.03.012

Douris, V., Denecke, S., Van Leeuwen, T., Bass, C., Nauen, R., Vontas, J., 2020. Using CRISPR/Cas9 genome modification to understand the genetic basis of insecticide resistance: *Drosophila* and beyond. Pesticide Biochemistry and Physiology 167, 104595. 10.1016/j.pestbp.2020.104595

EPPO, 2026. Tetanychus evansi [WWW Document]. EPPO datasheets on pests recommended for regulation. URL https://gd.eppo.int (accessed 2.18.26).

Fan, Q.-H., Li, D., Bennett, S., Balan, R.K., 2021. Diagnosis of a new to New Zealand spider mite, Tetranychus evansi Baker and Pritchard, 1960 (Acari: Tetranychidae). New Zealand Entomologist.

Feng, K., Ou, S., Zhang, P., Wen, X., Shi, L., Yang, Y., Hu, Y., Zhang, Y., Shen, G., Xu, Z., He, L., 2020. The cytochrome P450 CYP389C16 contributes to the cross-resistance between cyflumetofen and pyridaben in Tetranychus cinnabarinus (Boisduval). Pest Management Science 76, 665–675. 10.1002/ps.5564

Feyereisen, R., Dermauw, W., Van Leeuwen, T., 2015. Genotype to phenotype, the molecular and physiological dimensions of resistance in arthropods. Pesticide Biochemistry and Physiology, Insecticide and Acaricide Modes of Action and their Role in Resistance and its Management 121, 61–77. 10.1016/j.pestbp.2015.01.004

Field, L.M., Emyr Davies, T.G., O’Reilly, A.O., Williamson, M.S., Wallace, B.A., 2017. Voltage-gated sodium channels as targets for pyrethroid insecticides. Eur Biophys J 46, 675–679. 10.1007/s00249-016-1195-1

Fotoukkiaii, S.M., Wybouw, N., Kurlovs, A.H., Tsakireli, D., Pergantis, S.A., Clark, R.M., Vontas, J., Leeuwen, T.V., 2021. High-resolution genetic mapping reveals cis-regulatory and copy number variation in loci associated with cytochrome P450-mediated detoxification in a generalist arthropod pest. PLOS Genetics 17, e1009422. 10.1371/journal.pgen.1009422

Ghazy, N.A., Gotoh, T., Suzuki, T., 2019. Impact of global warming scenarios on life-history traits of Tetranychus evansi (Acari: Tetranychidae). BMC Ecol 19, 48. 10.1186/s12898-019-0264-6

González-Cabrera, J., Davies, T.G.E., Field, L.M., Kennedy, P.J., Williamson, M.S., 2013. An Amino Acid Substitution (L925V) Associated with Resistance to Pyrethroids in Varroa destructor. PLOS ONE 8, e82941. 10.1371/journal.pone.0082941

Grbić, M., Van Leeuwen, T., Clark, R.M., Rombauts, S., Rouzé, P., Grbić, V., Osborne, E.J., Dermauw, W., Thi Ngoc, P.C., Ortego, F., Hernández-Crespo, P., Diaz, I., Martinez, M., Navajas, M., Sucena, É., Magalhães, S., Nagy, L., Pace, R.M., Djuranović, S., Smagghe, G., Iga, M., Christiaens, O., Veenstra, J.A., Ewer, J., Villalobos, R.M., Hutter, J.L., Hudson, S.D., Velez, M., Yi, S.V., Zeng, J., Pires-daSilva, A., Roch, F., Cazaux, M., Navarro, M., Zhurov, V., Acevedo, G., Bjelica, A., Fawcett, J.A., Bonnet, E., Martens, C., Baele, G., Wissler, L., Sanchez-Rodriguez, A., Tirry, L., Blais, C., Demeestere, K., Henz, S.R., Gregory, T.R., Mathieu, J., Verdon, L., Farinelli, L., Schmutz, J., Lindquist, E., Feyereisen, R., Van de Peer, Y., 2011a. The genome of Tetranychus urticae reveals herbivorous pest adaptations. Nature 479, 487–492. 10.1038/nature10640

Grbić, M., Van Leeuwen, T., Clark, R.M., Rombauts, S., Rouzé, P., Grbić, V., Osborne, E.J., Dermauw, W., Thi Ngoc, P.C., Ortego, F., Hernández-Crespo, P., Diaz, I., Martinez, M., Navajas, M., Sucena, É., Magalhães, S., Nagy, L., Pace, R.M., Djuranović, S., Smagghe, G., Iga, M., Christiaens, O., Veenstra, J.A., Ewer, J., Villalobos, R.M., Hutter, J.L., Hudson, S.D., Velez, M., Yi, S.V., Zeng, J., Pires-daSilva, A., Roch, F., Cazaux, M., Navarro, M., Zhurov, V., Acevedo, G., Bjelica, A., Fawcett, J.A., Bonnet, E., Martens, C., Baele, G., Wissler, L., Sanchez-Rodriguez, A., Tirry, L., Blais, C., Demeestere, K., Henz, S.R., Gregory, T.R., Mathieu, J., Verdon, L., Farinelli, L., Schmutz, J., Lindquist, E., Feyereisen, R., Van de Peer, Y., 2011b. The genome of Tetranychus urticae reveals herbivorous pest adaptations. Nature 479, 487–492. 10.1038/nature10640

Guerra, F., De Rouck, S., Verhulst, E.C., 2026. SYNCAS-mediated CRISPR-Cas9 genome editing in the Jewel wasp, Nasonia vitripennis. Insect Molecular Biology 35, 48–55. 10.1111/imb.70002

Habel, J.C., Schmitt, T., 2018. Vanishing of the common species: Empty habitats and the role of genetic diversity. Biological Conservation 218, 211–216. 10.1016/j.biocon.2017.12.018

Heckel, D.G., 2014. Insect Detoxification and Sequestration Strategies, in: Annual Plant Reviews. John Wiley & Sons, Ltd, pp. 77–114. 10.1002/9781118829783.ch3

İnak, E., De Rouck, S., Demirci, B., Dermauw, W., Geibel, S., Van Leeuwen, T., 2024a. A novel target-site mutation (H146Q) outside the ubiquinone binding site of succinate dehydrogenase confers high levels of resistance to cyflumetofen and pyflubumide in *Tetranychus urticae*. Insect Biochemistry and Molecular Biology 170, 104127. 10.1016/j.ibmb.2024.104127

İnak, E., De Rouck, S., Koç-İnak, N., Erdem, E., Rüstemoğlu, M., Dermauw, W., Van Leeuwen, T., 2024b. Identification and CRISPR-Cas9 validation of a novel β-adrenergic-like octopamine receptor mutation associated with amitraz resistance in *Varroa destructor*. Pesticide Biochemistry and Physiology 204, 106080. 10.1016/j.pestbp.2024.106080

Ji, M., Vandenhole, M., De Beer, B., De Rouck, S., Villacis-Perez, E., Feyereisen, R., Clark, R.M., Van Leeuwen, T., 2023. A nuclear receptor HR96-related gene underlies large trans-driven differences in detoxification gene expression in a generalist herbivore. Nat Commun 14, 4990. 10.1038/s41467-023-40778-w

Jia, H., Peiling, L., Yuan, H., Wencai, L., Zhifeng, X., Lin, H., 2019. P8 nuclear receptor responds to acaricides exposure and regulates transcription of P450 enzyme in the two-spotted spider mite, *Tetranychus urticae*. Comparative Biochemistry and Physiology Part C: Toxicology & Pharmacology 224, 108561. 10.1016/j.cbpc.2019.108561

Khajehali, J., Van Nieuwenhuyse, P., Demaeght, P., Tirry, L., Van Leeuwen, T., 2011. Acaricide resistance and resistance mechanisms in Tetranychus urticae populations from rose greenhouses in the Netherlands. Pest Management Science 67, 1424–1433. 10.1002/ps.2191

Knihinicki, D., Gopurenko, D., Gillespie, P.S., Millynn, B., Dominiak, B., Rossiter, L., 2025. Detection of the invasive tomato red spider mite, Tetranychus evansi Baker & Pritchard (Acari: Prostigmata: Tetranychidae) in Australia based on morphological identification and DNA sequence analysis. 10.1101/2025.06.11.657326

Letunic, I., Bork, P., 2024. Interactive Tree of Life (iTOL) v6: recent updates to the phylogenetic tree display and annotation tool. Nucleic Acids Res 52, W78–W82. 10.1093/nar/gkae268

Li, J., Cai, W., Wang, L., He, J., Meng, Y., Liu, J., Xu, L., Feng, K., Shen, G., Wei, P., He, L., 2026. Functional analysis of inducible choline/carboxylesterase *CCE01* associated with fenpropathrin and abamectin detoxification in *Tetranychus urticae* (Koch). Pesticide Biochemistry and Physiology 219, 107003. 10.1016/j.pestbp.2026.107003

Li, X., Xu, Y., Zhang, H., Yin, H., Zhou, D., Sun, Y., Ma, L., Shen, B., Zhu, C., 2021. ReMOT Control Delivery of CRISPR-Cas9 Ribonucleoprotein Complex to Induce Germline Mutagenesis in the Disease Vector Mosquitoes Culex pipiens pallens (Diptera: Culicidae). J Med Entomol 58, 1202–1209. 10.1093/jme/tjab016

Lu, X., Vandenhole, M., Tsakireli, D., Pergantis, S.A., Vontas, J., Jonckheere, W., Van Leeuwen, T., 2023. Increased metabolism in combination with the novel cytochrome *b* target-site mutation L258F confers cross-resistance between the Qo inhibitors acequinocyl and bifenazate in *Tetranychus urticae*. Pesticide Biochemistry and Physiology 192, 105411. 10.1016/j.pestbp.2023.105411

Marcolungo, L., Vincenzi, L., Ballottari, M., Cecchin, M., Cosentino, E., Mignani, T., Limongi, A., Ferraris, I., Orlandi, M., Rossato, M., Delledonne, M., 2023. Structural Refinement by Direct Mapping Reveals Assembly Inconsistencies near Hi-C Junctions. Plants 12. 10.3390/plants12020320

Matsuda, N., Hinomoto, N., Daimon, T., 2025. Comparison of DIPA-CRISPR and SYNCAS gene editing methods in the predatory bug Orius strigicollis (Hemiptera: Anthocoridae). Appl Entomol Zool 60, 309–316. 10.1007/s13355-025-00923-x

McKenna, A., Hanna, M., Banks, E., Sivachenko, A., Cibulskis, K., Kernytsky, A., Garimella, K., Altshuler, D., Gabriel, S., Daly, M., DePristo, M.A., 2010. The Genome Analysis Toolkit: A MapReduce framework for analyzing next-generation DNA sequencing data. Genome Research 20, 1297–1303. 10.1101/gr.107524.110

Migeon, A., Dorkeld, F., 2026. A comprehensive database for the Tetranychidae [WWW Document]. Spider Mites Web Home. URL https://www1.montpellier.inrae.fr/CBGP/spmweb/public/ (accessed 1.29.26).

Migeon, A., Ferragut, F., Escudero-Colomar, L.A., Fiaboe, K., Knapp, M., de Moraes, G.J., Ueckermann, E., Navajas, M., 2009. Modelling the potential distribution of the invasive tomato red spider mite, Tetranychus evansi (Acari: Tetranychidae). Exp Appl Acarol 48, 199–212. 10.1007/s10493-008-9229-8

Millán-Leiva, A., Marín, Ó., De la Rúa, P., Muñoz, I., Tsagkarakou, A., Eversol, H., Christmon, K., vanEngelsdorp, D., González-Cabrera, J., 2021. Mutations associated with pyrethroid resistance in the honey bee parasite Varroa destructor evolved as a series of parallel and sequential events. J Pest Sci 94, 1505–1517. 10.1007/s10340-020-01321-8

Mocchetti, A., De Rouck, S., Naessens, S., Dermauw, W., Van Leeuwen, T., 2025a. SYNCAS based CRISPR-Cas9 gene editing in predatory mites, whiteflies and stinkbugs. Insect Biochemistry and Molecular Biology 177, 104232. 10.1016/j.ibmb.2024.104232

Mocchetti, A., Nikoloudi, A., Phuong, T.T.T., Jonckheere, W., Nguyen, D.T., De Clercq, P., Van Leeuwen, T., 2025b. Geographical distribution and incidence of pesticide resistance mutations in spider mite and thrips species from North Vietnam. Pesticide Biochemistry and Physiology 215, 106634. 10.1016/j.pestbp.2025.106634

Mocchetti, A., Nikoloudi, A.A., Vontas, J., De Rouck, S., Van Leeuwen, T., 2025c. CRISPR/Cas9 knock-out of nAChR α6 confers resistance to spinosyns in *Frankliniella occidentalis* and is associated with a higher fitness cost than target-site mutation G275E. Pesticide Biochemistry and Physiology 212, 106455. 10.1016/j.pestbp.2025.106455

Mocchetti, A., Steelant, P., Hosseinkhani, M., De Rouck, S., Khajehali, J., Van Leeuwen, T., 2026. Knockout of nAChR subunits in spider mites and their phytoseiid predators confers spinosyn cross-resistance and reveals a conserved mode of action in mites. Insect Biochemistry and Molecular Biology 104498. 10.1016/j.ibmb.2026.104498

Monjarás-Barrera, J.I., Sanchez-Peña, S.R., 2024. First record of the tomato red spider mite, Tetranychus evansi Baker and Pritchard (Acari: Tetranychidae) in Mexico, from cultivated and wild solanaceous plants. Acarologia 64, 164–171. 10.24349/78jc-6nev

Moran, N.A., Jarvik, T., 2010. Lateral Transfer of Genes from Fungi Underlies Carotenoid Production in Aphids. Science 328, 624–627. 10.1126/science.1187113

Nauen, R., Bass, C., Feyereisen, R., Vontas, J., 2022. The Role of Cytochrome P450s in Insect Toxicology and Resistance. Annual Review of Entomology 67, 105–124. 10.1146/annurev-ento-070621-061328

Navajas, M., de Moraes, G.J., Auger, P., Migeon, A., 2013. Review of the invasion of Tetranychus evansi: biology, colonization pathways, potential expansion and prospects for biological control. Exp Appl Acarol 59, 43–65. 10.1007/s10493-012-9590-5

Njiru, C., Saalwaechter, C., Mavridis, K., Vontas, J., Geibel, S., Wybouw, N., Van Leeuwen, T., 2023. The complex II resistance mutation H258Y in succinate dehydrogenase subunit B causes fitness penalties associated with mitochondrial respiratory deficiency. Pest Management Science 79, 4403–4413. 10.1002/ps.7640

Njiru, C., Xue, W., De Rouck, S., Alba, J.M., Kant, M.R., Chruszcz, M., Vanholme, B., Dermauw, W., Wybouw, N., Van Leeuwen, T., 2022. Intradiol ring cleavage dioxygenases from herbivorous spider mites as a new detoxification enzyme family in animals. BMC Biol 20, 131. 10.1186/s12915-022-01323-1

Nyoni, B.N., Gorman, K., Mzilahowa, T., Williamson, M.S., Navajas, M., Field, L.M., Bass, C., 2011. Pyrethroid resistance in the tomato red spider mite, Tetranychus evansi, is associated with mutation of the para-type sodium channel. Pest Management Science 67, 891–897. 10.1002/ps.2145

O’Reilly, A.O., Khambay, B.P.S., Williamson, M.S., Field, L.M., WAllace, B.A., Davies, T.G.E., 2006. Modelling insecticide-binding sites in the voltage-gated sodium channel. Biochem J 396, 255–263. 10.1042/BJ20051925

Rameshgar, F., Khajehali, J., Nauen, R., Bajda, S., Jonckheere, W., Dermauw, W., Van Leeuwen, T., 2019. Point mutations in the voltage-gated sodium channel gene associated with pyrethroid resistance in Iranian populations of the European red mite *Panonychus ulmi*. Pesticide Biochemistry and Physiology 157, 80–87. 10.1016/j.pestbp.2019.03.008

Riga, M., Myridakis, A., Tsakireli, D., Morou, E., Stephanou, E.G., Nauen, R., Van Leeuwen, T., Douris, V., Vontas, J., 2015. Functional characterization of the Tetranychus urticae CYP392A11, a cytochrome P450 that hydroxylates the METI acaricides cyenopyrafen and fenpyroximate. Insect Biochemistry and Molecular Biology 65, 91–99. 10.1016/j.ibmb.2015.09.004

Riga, M., Tsakireli, D., Ilias, A., Morou, E., Myridakis, A., Stephanou, E.G., Nauen, R., Dermauw, W., Van Leeuwen, T., Paine, M., Vontas, J., 2014. Abamectin is metabolized by CYP392A16, a cytochrome P450 associated with high levels of acaricide resistance in Tetranychus urticae. Insect Biochemistry and Molecular Biology 46, 43–53. 10.1016/j.ibmb.2014.01.006

Rinkevich, F.D., Du, Y., Dong, K., 2013. Diversity and convergence of sodium channel mutations involved in resistance to pyrethroids. Pesticide Biochemistry and Physiology, Special Issue: Advances in Vector and Urban Pest Management and Resistance 106, 93–100. 10.1016/j.pestbp.2013.02.007

Robertson, J.L., Jones, M.M., Olguin, E., Alberts, B., 2017. Bioassays with Arthropods, 3rd ed. CRC Press, Boca Raton. 10.1201/9781315373775

Sarmento, R.A., Lemos, F., Bleeker, P.M., Schuurink, R.C., Pallini, A., Oliveira, M.G.A., Lima, E.R., Kant, M., Sabelis, M.W., Janssen, A., 2011. A herbivore that manipulates plant defence. Ecology Letters 14, 229–236. 10.1111/j.1461-0248.2010.01575.x

Schwartz, D.C., Li, X., Hernandez, L.I., Ramnarain, S.P., Huff, E.J., Wang, Y.-K., 1993. Ordered Restriction Maps of Saccharomyces cerevisiae Chromosomes Constructed by Optical Mapping. Science 262, 110–114. 10.1126/science.8211116

Shirai, Y., Piulachs, M.-D., Belles, X., Daimon, T., 2022. DIPA-CRISPR is a simple and accessible method for insect gene editing. Cell Reports Methods 2. 10.1016/j.crmeth.2022.100215

Silva, P., 1954. Um novo ácaro nocivo ao tomateiro na Bahia. Boletim do Instituto Biologica da Bahia 1, 1–20.

Snoeck, S., Pavlidi, N., Pipini, D., Vontas, J., Dermauw, W., Van Leeuwen, T., 2019. Substrate specificity and promiscuity of horizontally transferred UDP-glycosyltransferases in the generalist herbivore *Tetranychus urticae*. Insect Biochemistry and Molecular Biology 109, 116–127. 10.1016/j.ibmb.2019.04.010

Tsakireli, D., Vandenhole, M., Spiros A., P., Riga, M., Balabanidou, V., De Rouck, S., Ray, J., Zimmer, C., Talmann, L., Van Leeuwen, T., Vontas, J., 2024. The cytochrome P450 subfamilies CYP392A and CYP392D are key players in acaricide metabolism in *Tetranychus urticae*. Pesticide Biochemistry and Physiology 204, 106031. 10.1016/j.pestbp.2024.106031

Udall, J.A., Dawe, R.K., 2018. Is It Ordered Correctly? Validating Genome Assemblies by Optical Mapping. Plant Cell 30, 7–14. 10.1105/tpc.17.00514

Van Leeuwen, T., Vanholme, B., Van Pottelberge, S., Van Nieuwenhuyse, P., Nauen, R., Tirry, L., Denholm, I., 2008. Mitochondrial heteroplasmy and the evolution of insecticide resistance: Non-Mendelian inheritance in action. Proceedings of the National Academy of Sciences 105, 5980–5985. 10.1073/pnas.0802224105

Winters, A.E., Stevens, M., Mitchell, C., Blomberg, S.P., Blount, J.D., 2014. Maternal effects and warning signal honesty in eggs and offspring of an aposematic ladybird beetle. Functional Ecology 28, 1187–1196. 10.1111/1365-2435.12266

Wong, T.K.F., Ly-Trong, N., Ren, H., Baños, H., Roger, A.J., Susko, E., Bielow, C., Maio, N.D., Goldman, N., Hahn, M.W., Huttley, G., Lanfear, R., Minh, B.Q., 2025. IQ-TREE 3: Phylogenomic Inference Software using Complex Evolutionary Models.

Wu, M., Adesanya, A.W., Morales, M.A., Walsh, D.B., Lavine, L.C., Lavine, M.D., Zhu, F., 2019. Multiple acaricide resistance and underlying mechanisms in Tetranychus urticae on hops. J Pest Sci 92, 543–555. 10.1007/s10340-018-1050-5

Wybouw, N., Dermauw, W., Tirry, L., Stevens, C., Grbić, M., Feyereisen, R., Van Leeuwen, T., 2014. A gene horizontally transferred from bacteria protects arthropods from host plant cyanide poisoning. eLife 3, e02365. 10.7554/eLife.02365

Wybouw, N., Kosterlitz, O., Kurlovs, A.H., Bajda, S., Greenhalgh, R., Snoeck, S., Bui, H., Bryon, A., Dermauw, W., Van Leeuwen, T., Clark, R.M., 2019a. Long-Term Population Studies Uncover the Genome Structure and Genetic Basis of Xenobiotic and Host Plant Adaptation in the Herbivore Tetranychus urticae. Genetics 211, 1409–1427. 10.1534/genetics.118.301803

Wybouw, N., Kosterlitz, O., Kurlovs, A.H., Bajda, S., Greenhalgh, R., Snoeck, S., Bui, H., Bryon, A., Dermauw, W., Van Leeuwen, T., Clark, R.M., 2019b. Long-Term Population Studies Uncover the Genome Structure and Genetic Basis of Xenobiotic and Host Plant Adaptation in the Herbivore Tetranychus urticae. Genetics 211, 1409–1427. 10.1534/genetics.118.301803

Xu, X., Harvey-Samuel, T., Yang, J., Alphey, L., You, M., 2020. Ommochrome pathway genes kynurenine 3-hydroxylase and cardinal participate in eye pigmentation in Plutella xylostella. BMC Mol and Cell Biol 21, 63. 10.1186/s12860-020-00308-8

Xue, W., Lu, X., Mavridis, K., Vontas, J., Jonckheere, W., Van Leeuwen, T., 2022. The H92R substitution in PSST is a reliable diagnostic biomarker for predicting resistance to mitochondrial electron transport inhibitors of complex I in European populations of Tetranychus urticae. Pest Management Science 78, 3644–3653. 10.1002/ps.7007

Xue, W.-X., Sun, J.-T., Witters, J., Vandenhole, M., Dermauw, W., Bajda, S.A., Simma, E.A., Wybouw, N., Villacis-Perez, E., Van Leeuwen, T., 2023. Incomplete reproductive barriers and genomic differentiation impact the spread of resistance mutations between green-and red-colour morphs of a cosmopolitan mite pest. Molecular Ecology 32, 4278–4297. 10.1111/mec.16994

Yan, W., Du, L., Liu, H., Li, G.-Y., 2024. Current and future invasion risk of tomato red spider mite under climate change. J Econ Entomol 117, 1385–1395. 10.1093/jee/toae140

Zhorov, B.S., Dong, K., 2017. Elucidation of pyrethroid and DDT receptor sites in the voltage-gated sodium channel. NeuroToxicology 60, 171–177. 10.1016/j.neuro.2016.08.013

